# Delivery of Peptide Coacervates to Form Stable Interaction Hubs in Cells

**DOI:** 10.1101/2024.11.26.625566

**Authors:** Wangjie Tu, Rachel Q. Theisen, Pengfei Jin, David M. Chenoweth, Amish J. Patel, Matthew C. Good

**Affiliations:** Bioengineering Graduate Group, University of Pennsylvania, PA 19104; Department of Cell and Developmental Biology, University of Pennsylvania, PA 19104; Chemistry Graduate Group, University of Pennsylvania, PA 19104; Department of Chemistry, University of Pennsylvania, PA 19104; Chemical and Biomolecular Engineering Department, University of Pennsylvania, PA 19104; Department of Bioengineering, University of Pennsylvania, PA 19104

## Abstract

Cells contain membrane-bound and membraneless organelles that operate as spatially distinct biochemical niches. However, these reaction centers lose fidelity due to aging or disease. A grand challenge for biomedicine is restoring or augmenting cellular functionalities. Although commonly tackled by gene replacement therapy, an exciting new strategy is the delivery of protein-based materials that directly interact with and alter intracellular pathways. In this study, we sought to develop long-lasting materials capable of cellular uptake, akin to artificial organelles or synthetic interaction hubs. We show efficient delivery of micron-size peptide-based compartments into a variety of cell types. By loading coacervates with nanobodies and bioPROTACs, we demonstrate successful native target sequestration, and hub bioreactor function to selectively degrade targets inside human cells. These results represent an important step toward the development of synthetic organelles that can be fabricated in vitro and taken up by cells for applications in cell engineering and regenerative medicine.

## Introduction

A grand challenge for biomedical science is achieving predictable control over cellular functions by controlling protein networks inside cells^1–3^. In nature, cells utilize spatial partitioning via anchoring and scaffold proteins and leverage subcellular compartmentalization as key strategies to enhance the efficiency of biochemical processes^4–6^. We previously demonstrated the feasibility of controlling cellular processes and cell behavior by expressing proteins capable of forming synthetic membraneless organelles intracellularly and targeting endogenous enzymes^7,8^. However, to date this approach has required gene delivery using viruses or nanoparticles. A major advance would be the ability to fabricate compartments ex vivo and deliver them into cells to stably function as designer synthetic hubs or organelles.

Cells control the rate and fidelity of biochemical reactions by compartmentalizing components in membrane-bound and membraneless organelles. For example, oxidative phosphorylation is caried out in lipid bilayer containing mitochondria^9^, and membraneless organelles such as the nucleolus specifically govern rRNA processing^10–12^. Stress granules sequester factors to downregulate protein translation^13^ and Nck-N-WASP condensates^14,15^ recruit Sos1 and Arp2/3 to to form a reaction hub^16^. Membraneless compartments are formed by the self-assembly of proteins and RNAs intro micron-size condensates driven via multivalent interactions^6,17,18^. Similar ideology can be adopted to generate new designer condensates or compartments to programmably control reaction pathways in a cell^7,19,20^. Often the proteins that participate in this self-assembly and condensation contain large disordered or low-complexity polypeptide sequences^6,21^. We previously created synthetic condensates by expressing a disordered scaffold protein in cells and used them to reprogram cell behaviors. For example, we built compartments that functioned as insulators, turning signal flow on or off, to regulate growth and division or to polarize cytoskeleton organization^7^. Separately, synthetic condensates can be used as bioreactors to enhance signaling between components or generate new signaling cascades among factors that normally only weakly interact^20,22^. However, there are limitations to the use of these approaches for clinical applications: gene delivery is required, gene expression has a maximum^23^ that limits the amount of condensed material (expression relative to c_sat_)^8^, and native targets must be tagged to drive their localization to the synthetic condensate^7,8^. We sought to overcome these hurdles using a strategy for protein-based delivery via peptide coacervates.

There are a variety of strategies to achieve protein or nucleic acid delivery to cells. Commonly the RNA for a target of interest is delivered via lipid nanoparticles, which then drives expression inside a cell^24^. However it is not feasible to achieve very high protein expression with these methods^23,25^, which limits their utility for delivering materials to form protein condensates that function as synthetic organelles. Cell permeable peptides exist in nature and have been designed to be capable of crossing the plasma membrane^26–28^. Although useful for drug delivery, these short peptide sequences cannot simply be appended to target proteins of interest to drive their cellular uptake. Recently, a promising strategy for protein delivery has emerged based on identification of peptide coacervate particles capable of crossing the plasma membrane for cellular uptake^29,30^. A 26 amino acid disordered peptide HBpep was identified and shown capable of self-assembly into micron size coacervates through liquid-liquid phase separation^29,31,32^. These particles readily cross the cell membrane for cytosolic delivery. These coacervates can be co-assembled or loaded with cargos including proteins and RNAs and will partially disassemble upon entering the cells, highlighting their utility for efficiency cytosolic delivery of macromolecules^29,33,33,34^. Variants of HBpep were developed such as HBpep-SA and HBpep-SP, which are sensitive to redox state and partially disassemble in the cytosol of a cell^29^. Although primarily designed for drug delivery, we reason that the HBpep platform could be adapted to create stable protein-based hubs or organelles in cells, creating a toolbox to augment or revitalize cellular functions.

In this study, we create a platform to form large peptide coacervates capable of uptake and stable maintenance inside living cells. We develop strategies to enhance protein cargo loading and topological organization in coacervates, moving beyond passive loading and uniform partitioning. In contrast to previous controlled release systems, we develop strategies to deliver micron-size coacervates to cells that can serve as synthetic hubs or compartments, stable for days. Further we embed nanobodies within these coacervates and demonstrate them capable of interfacing with protein targets in vitro and in cells. Finally, by co-loading bioPROTACs, we demonstrate that these deliverable intracellular coacervates can function as reaction centers for degradation. Delivery of synthetic hubs capable of targeting native enzymes and signaling pathways provides a powerful strategy in cellular engineering. By eliminating the requirement to deliver genetic material, this modular platform promises to fast-track therapeutic development, for example through formation of degradomes or via compartments that rewire aberrant signaling responses.

## Results

### Parameter optimization to form stable micron-size protein coacervates

Building from a previous peptide coacervate platform, HBpep^31,33^, we sought to develop a protocol to form gel-like particles that range in size from 1-5 µm diameter for cargo loading and cellular uptake, and that could ultimately serve as stable synthetic hubs or organelles in a cell. We chose to promote nucleation phase separation using two different methods: pH shift and temperature shift. HBpep contains five histidines and is largely soluble in acidic buffer, such as pH 4.5, but will phase separate at physiological pH 7.4 under standard conditions^35^. Further, HBpep displays upper critical solution temperature behavior and LLPS can be induced by shifting the temperature from 50°C to room temperature, 23°C. We also chose to test both the standard HBpep, which has only limited cargo release capability and a redox sensitive form HBpep-SA, that will release cargo into the cytoplasm of a cell^29^.

We first measured coacervation via pH shift for HBpep and HBpep-SA over a range of peptide concentrations from 0.1 to 2 mg/mL, corresponding to approximately 35 – 700 micromolar. We found coacervate size and number depended on peptide concentration (Fig. 1A-C), as expected^36^. Although smaller particles have utility for drug delivery applications, we were interested in formation of larger hubs that would be stably maintained in cells. At concentrations of 0.5 mg/mL or higher, the average HBpep particle diameter was larger than 2 µm and could range from 1-6 µm (Fig. 1B). Similar results were attained with HBpep-SA, although the sizes were slightly smaller. A higher fraction of the HBpep-SA peptide partitioned to the coacervates as measured by sedimentation of particles (Supplementary Fig1A-B), although in both cases at majority of the scaffold peptide was condensed when assembled at a concentration of 0.5 mg/mL. We separately tested induction of LLPS via temperature shift and observed similar size and numbers of particles compared to induction via pH shift (Supplementary Fig. 1C-E). During protocol optimization, we found that the free peptides in solutions at pH 7.4 were actually quite sticky and would foul surfaces of test tubes or imaging chambers over time. Particle production via pH shift is somewhat more reliable at creating dose-dependent coacervation and indicates use of acidic buffers for storage and handling of HBpep and HBpep-SA prior to assembly.

**Figure 1.**
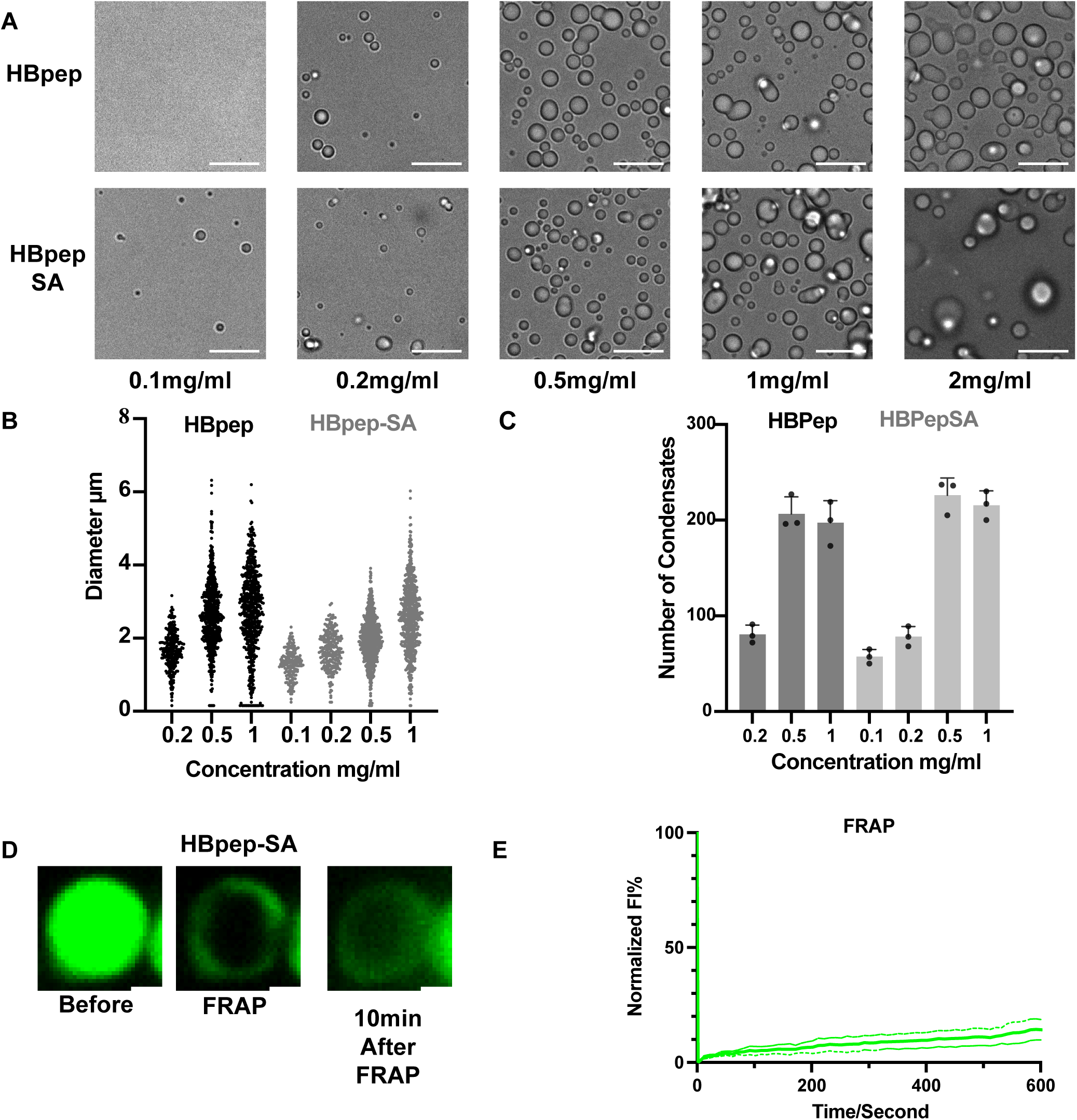
Assembly and size control of HBpep and HBpep-SA gel-like peptide coacervates. **(A)** Images showing assembly of coacervate particles from HBpep and HBpep-SA using pH shift method at different concentrations of peptide. Solutions were at room temperature, 100 mM NaCl and at pH 7.4. Scale bar: 10 µm. **(B)** Quantification of HBpep and HBpep-SA coacervate diameters at different concentrations via pH shift method. The number of coacervates used for quantification per group, n=242, 620, 592, 172, 235, 678, 647 from left to right groups. **(C)** Quantification of average number of HBpep and HBpep-SA coacervates upon assembly at different concentrations (number per 4616.2 µm^2^ imaging area). The number of 4616.2 µm^2^ images used per group, n=3. Error bars indicate standard deviation from mean. Center of error bars indicate the mean. **(D)** Fluorescence images over time of FITC labeled (1:100) HBpep-SA coacervates formed at 0.5 mg/mL via pH shift, following laser photobleaching. Scale bar: 1 µm. **(E)** Quantification of normalized fluorescence after photobleaching, showing intensity within bleached region of FITC labeled HBpep-SA coacervates formed via pH shift. Source data are provided as a Source Data file.

For coacervates to be useful for non-genetic delivery of ‘hubs’ to cells, they need to show gel-like material properties and not require free peptide present in the continuous phase to maintain their assembly. In contrast, liquid-like condensates of disordered proteins formed inside cells from gene expression can have high turnover because the protein is continuously expressed and maintained by a pool of material in the cytosol^37^. We thus tested the extent to which coacervates were in a gel-like or more liquid-like state by measuring labeled peptide diffusion following fluorescence recovery after photobleaching (FRAP). Particles were co-assembled with tracer amounts of a FITC-labeled HBpep (FITC-HBpep) (Supplementary Fig. 1F) peptides and bleached using a 405 nm laser. Fluorescence recovery was quite limited over time, for both types of peptides, indicating very slow diffusion and suggesting that these particles are gels (Fig. 1D,E), as expected^34^. Recovery was slow but detectable over 10 minutes when photobleaching a portion of the particle to measure diffusion internally within the coacervate (Supplementary Fig. 1G, H). Fluorescence recovery from full photobleaching of particles was nearly undetectable, suggesting minimal recovery via diffusion from the continuous phase. We conclude it is feasible to generate particles of 1-5 µm in diameter using either method to induce LLPS and that peptide concentration can be varied to tune average coacervate size. Given their gel-phase properties we expected it would be feasible to dilute them into new solutions without providing additional material in continuous phase and that they would remain stable.

### Enhanced cargo loading into peptide coacervates via selective partitioning

We wondered whether it would be feasible to enhance selectivity and overall amount of cargo loaded into coacervates by tagging the cargo. Previous studies demonstrated non-specific entrapment or loading of cargos such as protein and RNA during assembly of HBpep and HBpep-SA coacervates^29^. However, loading efficiency varies for different cargos and depends on HBpep peptide concentration^29^. In this study, we explored increasing the specificity of cargo loading without having to adapt the cargo or change HBpep peptide concentration. We used GFP as a model folded protein cargo for this proof-of-concept test. To promote selective loading, we added a 25-amino-acid motif from HBpep (lacking tryptophan): GHGVY GHGVY GHGPY GHGPY GHGLY to the C-terminus of GFP, which we term GFP-HBP (Fig. 2A), and we generated the protein via recombinant expression and purification. We co-assembled the cargo via LLPS of HBpep and measured extent of loading. We compared loading of GFP vs GFP-HBP both for HBpep and HBpep-SA particles, in which particles were formed by pH shift. We found that HBpep enhanced cargo loading into coacervates (Fig. 2B-D), without changing coacervate sizes. We observed a nearly 2- to 3-fold increase in internal concentration of cargo as measured by fluorescence microscopy (Fig. 2C, D) and total partitioning as measured via sedimentation (Supplementary Fig. 2A, B). We conclude it is feasible to enhance the overall magnitude of loading and internal concentration of cargo by fusion of an HBP tag to a protein of interest. Under conditions in which cargos would compete, tagging would provide enhanced selectivity.

**Figure 2.**
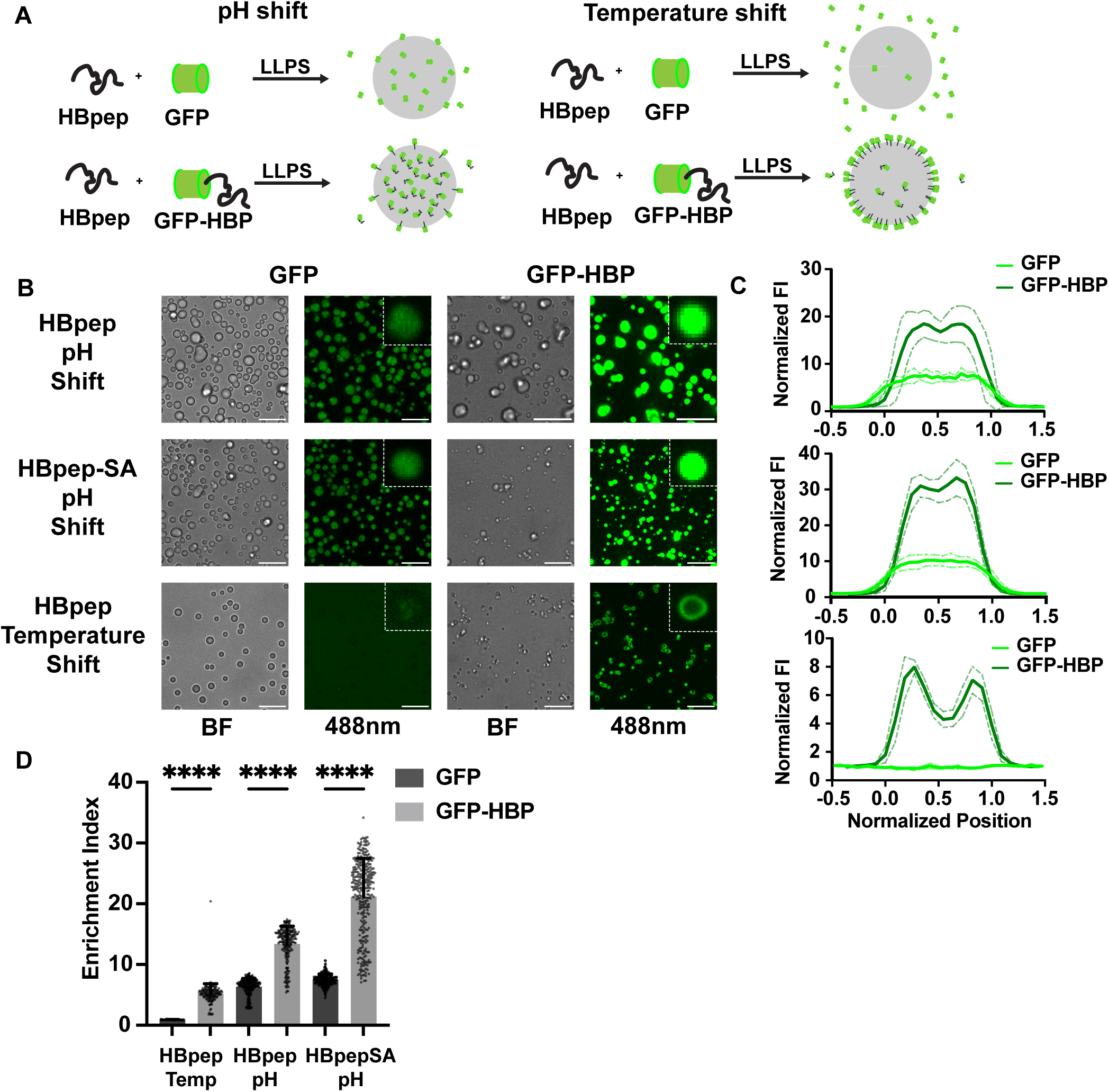
Enhanced cargo loading and topological partitioning using disordered peptide tags. **(A)** Scheme showing strategy to enhance selectivity of cargo loading: comparing of GFP vs tagged GFP-HBP loaded in HBpep condensates through pH shift (left) or temperature shift (right) method. **(B)** Images of cargo loading into coacervates: 0.1 mg/mL GFP or GFP-HBP co-assembled (loaded) into 0.5mg/mL HBpep or HBpep-SA particles through pH or temperature shift. Scale bar: 10 µm. **(C)** Quantitation of cargo loading via normalized line scan, averaging GFP or GFPHBP in coacervates of similar size. n=5 from (B). **(D)** Quantification of fluorescence intensity of cargo loading for GFP and GFPHBP in HBpep or HBpep-SA coacervates. Enrichment index: fluorescence intensity in coacervate divided by the continuous phase. The number of coacervates used for quantification per group, n=67,149, 292, 227, 321, 321 from left to right groups. Error bars indicate the standard deviation from mean. Center of error bars indicate the mean. Brown-Forsythe and Welch One-Way ANOVA tests were used. Dunnett T3 statistical hypothesis is used to correct for multiple comparisons. NS indicates not significant (P > 0.05), *P ≤ 0.05, **P ≤ 0.01, ***P ≤ 0.001 and ****P ≤ 0.0001. P-value ≤ 0.0001 in these 3 comparisons. Source data are provided as a Source Data file.

We next wondered whether it would be feasible to control the topology of cargo loading, dictating its spatial partitioning in the coacervate. Cargo loading appears relatively spatially uniform for coacervates formed via pH shift method, and in general cargos reach a higher internal concentration in particles comparing LLPS of HBpep via pH versus temperature for HBpep (Fig. 2B). Tagging greatly increases the partitioning of GFP to particles formed via temperature shift (Fig. 2C). Interestingly, the HBP tag also dictates the topology of cargo partitioning. GFP alone as a cargo shows relatively spatial uniform partitioning within particles, whereas GFP-HBP enriches at the particle surface (Fig. 2B, C). We conclude it is possible to specify a core-shell organization of scaffold and cargo that would enable spatially partitioning discrete cargos and would be broadly useful for a variety of applications in drug delivery and cell engineering. Nonetheless, for the remainder of this study we utilize pH shift to generate particles with uniform loading.

### Delivering micron size or larger coacervates to cells and characterizing their stability

A previous study focused on delivery of particles smaller than 1 µm to cells and for the purpose of cytosolic particle dissociation and cargo release^29^. However, in this study, we were interested in using the HBpep and HBpep-SA platforms as stable synthetic hubs or organelles inside cells. As a result, we wanted to test the feasibility of delivering larger condensates to cells and their stability following uptake inside cells.

We sought to identify conditions for highly efficient uptake of large particles in a variety of cell types. This included a transformed human cell line for osteosarcoma, U2OS, a mouse melanoma cell line, B16-F10, and primary human immune cells - monocytes (Fig. 3A). We pre-assembled particles at 0.5 mg/mL, diluted them in cell media from 1:10 to 1:3 and measured particle uptake after 4 hrs and 24 hrs. Cells were washed extensively after particle addition to remove those not delivered to cells. We observed generally highly efficient uptake by cells for both particles of HBpep and HBpep-SA, in a variety of cell types (Fig. 3A). These particles were present in the cytosol, but not the nucleus (Fig. 3A) and showed diffusive motion (*Supp Movie1*). A recent study suggests these coacervates cross the cell membrane via macropinocytosis which enables the uptake of large portions of extracellular material^32^. At 1 day, we measured effective and robust uptake of particles formed at both 0.2 and 2 mg/mL, averaging uptake of nearly 40 particles per cell at higher concentrations (Fig. 3B-E). Particle formulations include a range of sizes although those formed at 0.2 mg/mL, the mode for the distribution of diameters was less than 1 µm and those formed at 2 mg/mL were generally larger than 1 µm. Importantly there was a long tail in the distribution of particle sizes at higher concentrations (2 mg/mL formulation), and cells given these formulations contained nearly 10-fold higher numbers of larger particles, greater than 2 µm diameter (Fig. 3D, E). In exemplary cases we found it possible to fill up the cytoplasm of a cell with approximately 100 particles (Supplementary Fig. 3E, *Supp Movie 2*), a feat not possible from genetic expression of a coacervating protein. Therefore, it is feasible for larger coacervates to be efficiently taken up by cells.

**Figure 3.**
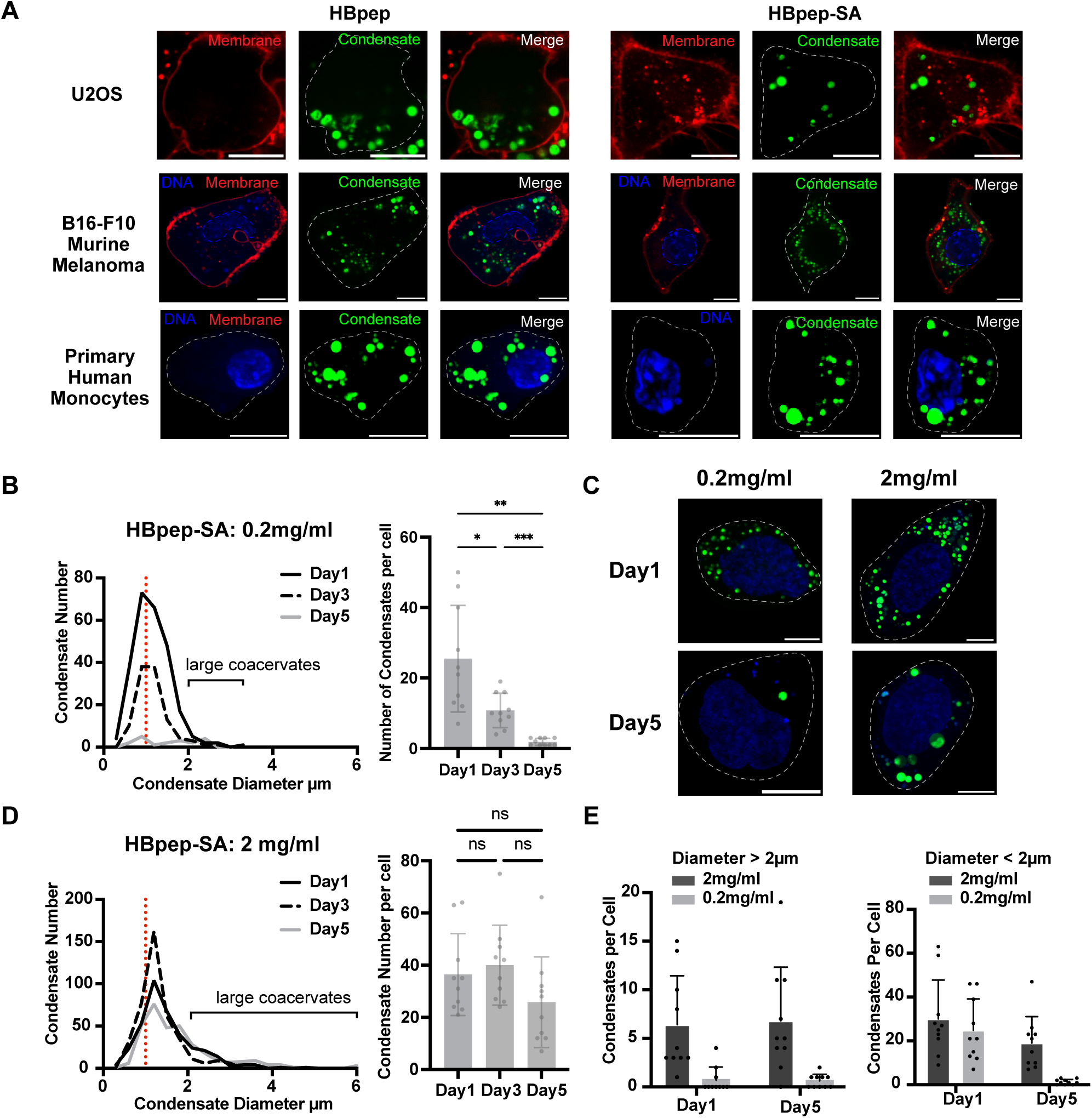
Stable delivery of peptide coacervates to mouse and human cells and clinically relevant cell types. **(A)** Fluorescence images of coacervate uptake in cell lines. 30 µl of HBpep particles assembled at 1mg/mL or 30 µl HBpep-SA particles assembled at 0.5mg/mL were added to the cell media for human osteosarcoma line U2OS, B16-F10 murine melanoma cells, and primary human monocytes. Green: FITC labeled condensates. Red: Abcam Cytopainter Cell Membrane Staining. Blue: Nucleus, stained DNA (DAPI). Scale bar: 10 µm. **(B)(D)** Quantification of size and number of coacervate taken up per cells; for formulations of 20 µl 0.2 mg/mL or 10 µl 2 mg/mL HBpep-SA delivered to U2OS; following particles over the course of 5 days. For coacervate size quantification, n=255, 108, 19 for 0.2mg/mL from Day1 to Day5 and n= 364, 400, 258 for 2mg/mL from Day1 to Day5. For coacervate number per cell quantification, cell number n=10. Error bars indicate standard deviation from mean. Center of error bars indicate the mean. Brown-Forsythe and Welch One-Way ANOVA tests were used. Dunnett T3 statistical hypothesis is used to correct for multiple comparisons. NS indicates not significant (P > 0.05), *P ≤ 0.05, **P ≤ 0.01, ***P ≤ 0.001 and ****P ≤ 0.0001. Adjusted p value is 0.0389, 0.0006, 0.0024 for B and 0.9365, 0.1850, 0.4135 for D. **(C)** Fluorescence images of U2OS cells with particles at day 1 and day 5. Scale bar: 10 µm. **(E)** Quantification of the number of large coacervates (diameter > 2 µm) per cell compared to smaller ones (diameter < 2 µm) for particles assembled at 2 mg/mL and 0.2 mg/mL HBpep-SA. The number of cells used for quantification per group, n=10. Error bars indicate the standard deviation from mean. Center of error bars indicate the mean. NS indicates not significant (P > 0.05), *P ≤ 0.05, **P ≤ 0.01, ***P ≤ 0.001 and ****P ≤ 0.0001. Source data are provided as a Source Data file.

In an effort to build stable hubs that can be delivered to cells, we were interested in the stability of the particles inside cells over time. Previous efforts focused on use of particles < 1 µm diameter for drug delivery that would subsequently partially disassemble in the cytoplasm^29^. We sought to test the stability of the intracellular coacervates, comparing formulations better suited for drug delivery versus those ideal for making stable hubs (Supplementary Fig. 3A-B). Particles were generally more stable at day 5 for coacervates assembled at 2 mg/mL vs. 0.2 mg/mL, likely related to their larger sizes (Fig. 3B, D and Supplementary Fig. 3C). Indeed, if when we quantified only larger particles (those > 2 µm diameter), the number per cell was relatively constant over time (Fig. 3E). In contrast smaller particles (< 2 µm diameter) were lost from cells for the 0.2 mg/mL formulation. We suspect the disassembly is caused by protonated or reduced HBpep-SA peptides diffusing away. This whole process is kinetically slow especially for large coacervates, making larger coacervates more “stable”. We conclude that our large coacervates are efficiently taken up by cells, and that these particles are stable intracellularly for at least 5 days. These results provide a roadmap toward delivery and maintenance of synthetic ‘hubs’ within living cells without the need for gene transfer.

### Selective targeting of proteins to pre-formed coacervates using nanobodies in vitro

To be useful as a designer organelle or hub, coacervates must be functionalized with molecules that enable them to interface with native protein interaction networks or signaling pathways. Native membraneless organelles can function as insulators^13^ – sequestering components – or as bioreactor compartments that co-localize components at elevated concentrations^15^. We previously demonstrated that synthetic condensates could be formed via expression of a scaffold protein in cells, and that they could function as insulators, targeting and partitioning native enzymes that were tagged with a complementary coiled coil^7^. In this study we wanted to go further: to deliver synthetic hubs via extracellular uptake rather than via gene transfer and expression, and to embed nanobodies within them that could bind to a target of interest without needing to tag the target. It would then be possible to create a designer biochemical niche inside a cell through the delivery of a one-pot peptide coacervate platform.

To start, we tested the feasibility of targeting proteins to pre-assembled peptide coacervates in vitro comparing those containing or lacking a nanobody (Fig. 4A). As a proof-of-concept we tested co-assembly and embedding of the nanobody VHHGFP4 which binds with nanomolar affinity to GFP^38^. This nanobody is highly useful because many stable cell lines have been created that contain a GFP-fusion to a target protein of interest. To track nanobody loading using fluorescence microscopy we fused it to mCherry (VHHGFP4-mCherry), or also to the HBP targeting sequence (VHHGFP4-mCherry-HBP). To form particles, these purified proteins were added a 1:5 mass ratio into formulations of 0.5mg/mL HBpep and HBpep-SA. Both nanobody cargos enriched within the HBpep and HBpep-SA particles after co-assembly (Supplementary Fig. 4A). We also observed somewhat elevated loading of VHHGFP4-mCherry-HBP into HBpep particles (Supplementary Fig. 4B), consistent with the results described above in Figure 2, that HBP-tagging of proteins enhances cargo loading during co-assembly with HBpep or HBpep-SA peptide.

**Figure 4.**
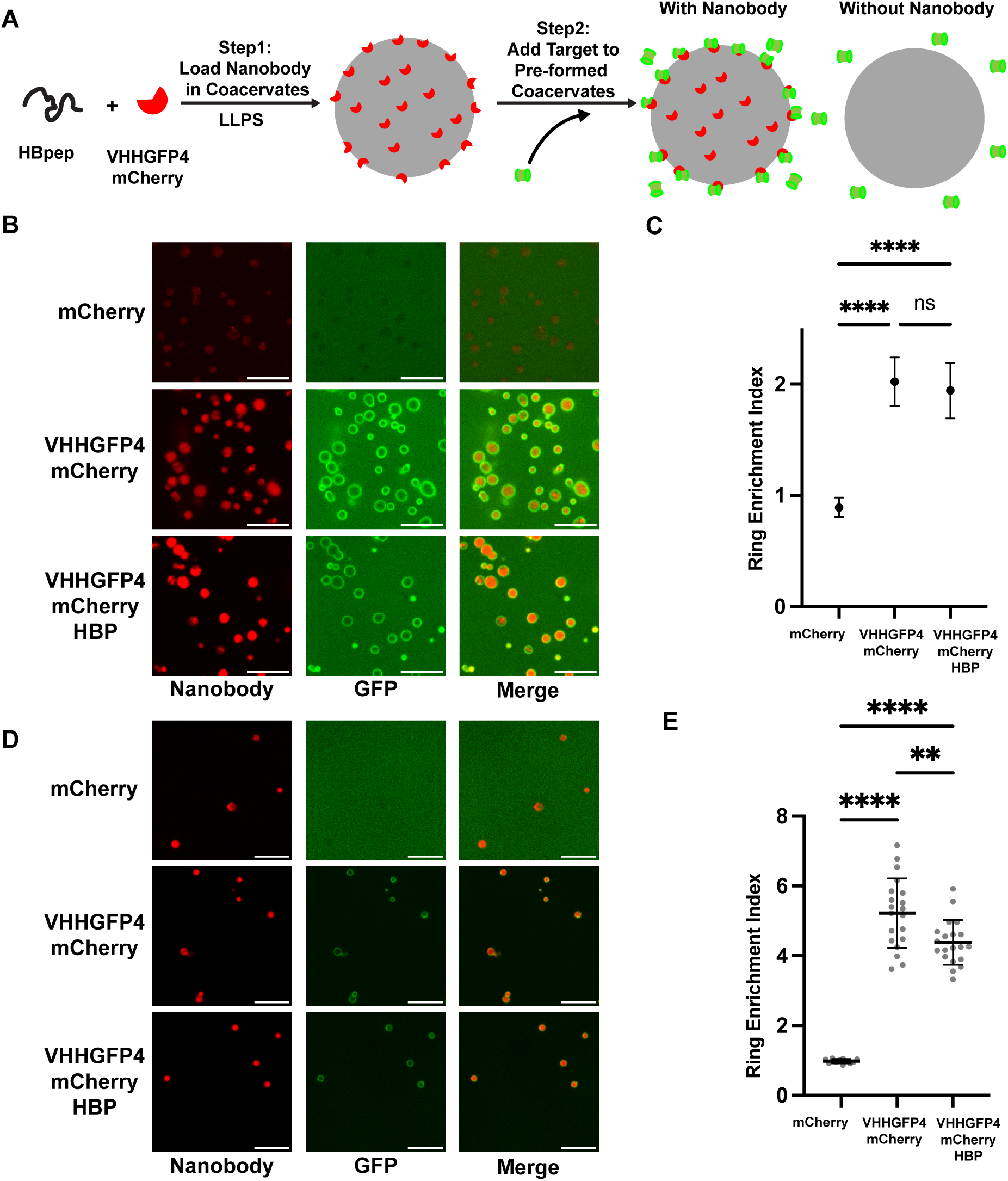
Selective binding of protein targets to nanobody containing coacervates in vitro. **(A)** Scheme to determine selectivity of target binding to preformed coacervates containing mCherry or VHHGFP4-mCherry. VHHGFP4 nanobody binds GFP. **(B)** Representative fluorescence images showing (i) cargo loading for Cherry or VHHGFP4 mCherry, and (ii) target protein (GFP) binding to coacervates following addition of 1 µM target protein. Images taken 5 min after GFP addition. scale bar: 10 µm. **(C)** Quantification of the enrichment of target GFP to HBpep-SA coacervates. Ring Enrichment Index: intensity of fluorescence on particle surface division by surrounding solution.: n=20 per group. Error bars indicate standard deviation from the mean. Center of error bars indicate the mean. Brown-Forsythe and Welch One-Way ANOVA tests were used. Dunnett T3 statistical hypothesis is used to correct for multiple comparisons. NS indicates not significant (P > 0.05), *P ≤ 0.05, **P ≤ 0.01, ***P ≤ 0.001 and ****P ≤ 0.0001. Adjusted p-value is P ≤ 0.0001 for two comparisons and 0.6401 for the ns one. **(D)** Representative fluorescence images showing cargo loading (mCherry or VHHGFP4 mCherry) and target protein (GFP) recruitment to hubs after dilution of free nanobody (1:10 dilution of coacervates in PBS prior to addition of <u>1uM of</u> GFP protein). Images taken 5 min after GFP addition. scale bar: 10 µm. **(E)** Quantification of the enrichment of target GFP to diluted HBpep-SA coacervates. Error bars: standard deviation from the mean. Center of error bars indicate the mean. n=11 for mCherry and n=20 for VHHGFP4mCherry. Brown-Forsythe and Welch One-Way ANOVA tests were used. Dunnett T3 statistical hypothesis is used to correct for multiple comparisons. Statistics: NS indicates not significant (P > 0.05), *P ≤ 0.05, **P ≤ 0.01, ***P ≤ 0.001 and ****P ≤ 0.0001. Adjusted p-value is P ≤ 0.0001 for two comparisons and 0.0090 for the **. Source data are provided as a Source Data file.

Once particles were formed, we quantitatively measured partitioning of GFP target protein added to pre-formed coacervates in an imaging microwell. Particles lacking a VHHGFP4 nanobody did not enrich GFP and instead partially excluded it (Fig. 4B, C); note partition coefficient of less than 1. Thus, cargo uptake is almost non-existent following peptide coacervate formation. In contrast particles containing VHHGFP4 nanobody enriched the GFP target at their interface (Fig. 4B) and this enrichment or partition coefficient at the surface was approximately 2-fold (Fig. 4C, Supplementary Fig. 4C, D) dependent on the type of peptide and whether HBP was present or not on the nanobody. This enrichment could be further enhanced to 4-to 6-fold by diluting the particles (Fig. 4D, E), which is consistent with excess nanobody free in solution competing with recruitment to the hubs. Note that the nanobody loading is relatively uniform throughout particles, and thus the surface targeting is not caused by spatial partitioning of nanobody to periphery. It was recently shown that these gel-like particles of HBpep and HBpep-SA have limited permeability for macromolecules larger than 2-10 kDa^39^. At 27 kDa in molecular weight, the target GFP would be expected to have limited diffusion inside the coacervate in the time course of our experiments, consistent with the ring-like GFP partitioning to our VHHGFP4 containing coacervates (Fig. 4B, D, Supplementary Fig. 4C)

Although mostly surface recruitment was achieved for this in vitro test using VHHGFP4 and GFP, it should be possible to achieve higher internal partitioning of molecules with smaller molecular weights, or by increasing the porosity of the coacervates. Notably for drug delivery, the cargo can be co-assembled with the particle as we show in Figure 2. To conclude, in this initial test we sought and found proof of feasibility for selective binding of target proteins to coacervates using nanobodies. Next, we wanted to test these particles for targeting in living cells.

### Target enrichment at synthetic protein hubs inside cells

It has been suggested that HBpep particles are taken up by cells through actin-mediated macropinocytosis and phagocytosis^32^. In such a process, the coacervates will be encapsulated by the endosomal lipid membrane, which limits accessibility and thus binding of target protein proteins. Based on published results and our own GFP delivery experiments using HBpep-SA particles (Supplementary Fig. 3D), some GFP can be released into the cytosol, which implies that there may be some stochastic escape from membrane encapsulation. However to further promote endosomal escape, we included TAT-HA2 assist peptide^40^ in co-assembly of HBpep-SA particles loaded with VHHGFP4 nanobody or a control protein. This TAT-HA2 assist peptide has been shown to create nanopores in the endosomal membranes that promote endosomal escape^26,40,41^. As a test case, we used a U2OS human cell line that was virally transduced to stably express GFP as a target protein.

To determine whether coacervates could function as hubs in cells we wanted to compare target GFP binding to control mCherry containing HBpep-SA particles versus those containing the VHHGFP4-mCherry nanobody. In cells loaded with control particles, we observed that most coacervates excluded the target GFP (Fig. 5A, C), showing a partition coefficient less than 1. In contrast for VHHGFP4-mCherry group, although many coacervates did not sequester the target (likely due to presence of membrane), we saw a large significant jump in target binding (Fig. 5B, D). The partition coefficients achieved both for scaffold and target are on par with our previous study which used coiled-coil recruitment of targets into RGG condensates^7^. In cells, enrichment indices below 2 are often non-specific. For control mCherry particles target GFP enrichment was below this value in nearly every case; we observed it < 1% of time (Fig. 5E). In contrast hubs containing VHHGFP4-mCherry and TAT-HA2 assist peptide showed target GFP partitioning as high as 15-fold above cytosolic levels (Fig. 5D, Supplementary Fig. 5A). The percentage of condensates with GFP enrichment > 2 was approximately 11.5% for particles co-assembled with VHHGFP4-mCherry and TAT-HA2, representing an order of magnitude increase in target enrichment compared to control particles, and comparable to the rates of efficiency of endosomal escape for some LNPs^42,43^. There is also a general trend of increasing average GFP enrichment among condensates with GFP enrichment > 1 with VHHGFP4-mCherry and TAT-HA2 loaded compared to control group (Supplementary Fig. 5B). At a single-cell level, the fraction of cells that show at least one hub with significant target recruitment is 7-10% in control hubs and as high as 47-60% in hubs co-assembled with the VHHGFP4-mchery nanobody (Fig. 5F). These results demonstrated initial feasibility of the platform and also hinted that additional refinements were needed to control or enhance the target recruitment process.

**Figure 5.**
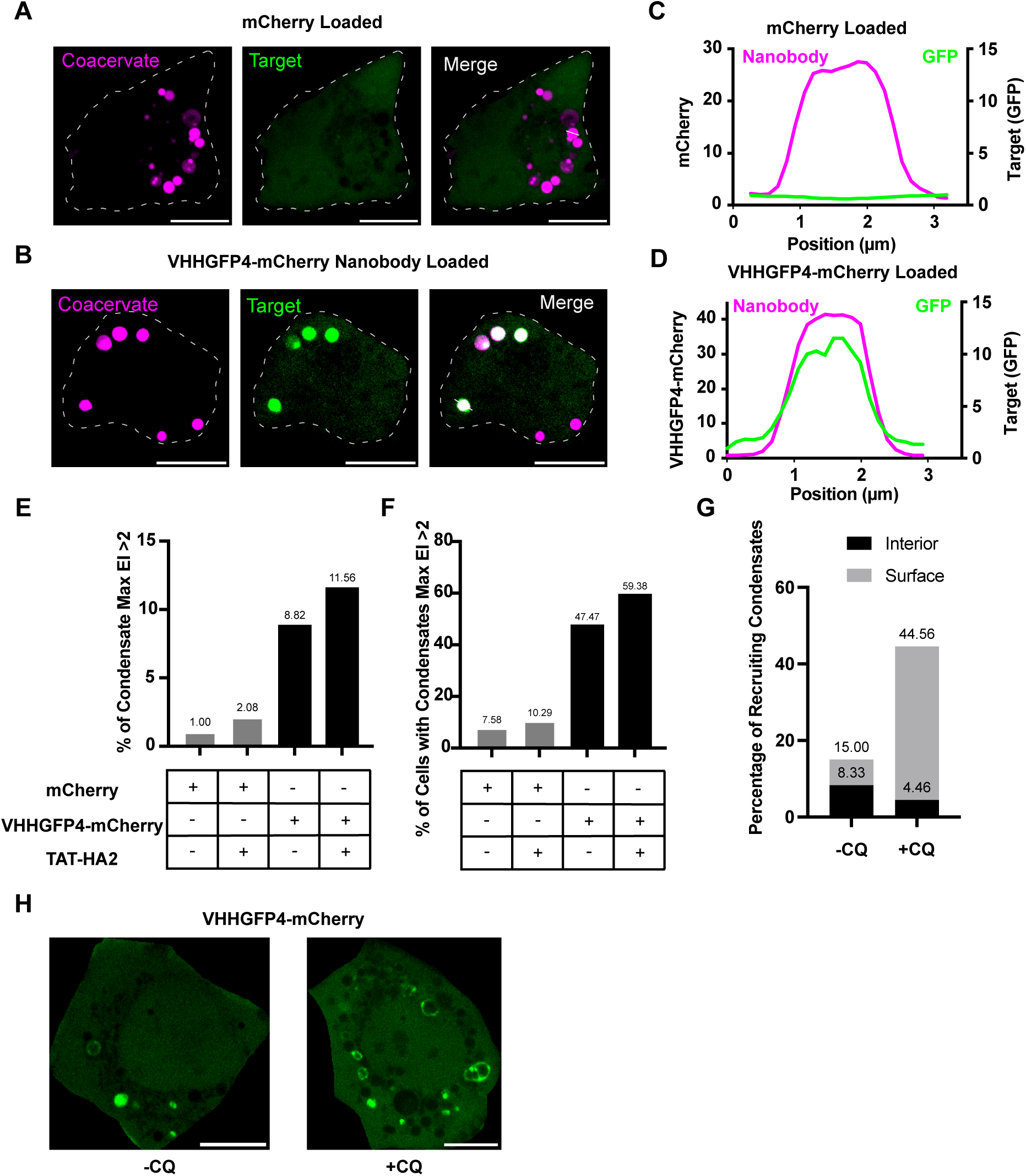
Delivered VHHGFP4 HBpep-SA “Hubs” bind to and enrich native target GFP protein. **(A)(B)** Representative fluorescence images of target binding to hubs inside U2OS cells stably expressing GFP, comparing mCherry control and VHHGFP4(nanobody)mCherry-loaded coacervates. 30µl of coacervates formed from 0.5 mg/mL HBpep-SA, co-assembled with 0.1mg/mL mCherry or VHHGFP4-mCherry were added to the cell media for U2OS cells and imaged at 24 hr. Cell boundaries are indicated by dashed gray line. Chosen ‘hub’ for line scan is indicated by white line. Scale bar: 10 µm. **(C)(D)** Line scans of coacervate hubs showing fluorescence enrichment in (A)(B). **(E)** Quantification: percentage of hubs in cells that show max enrichment index greater than 2. Sample size n= 902, 865, 1145, 986. **(F)** Quantitation of percentage of cells that show at least one hub recruiting target with EI >2. N=66, 68, 99, 96. **(G)** Quantitation of percentage of hubs that show either average interior enrichment index >2 or average surface enrichment index >1.1. n= 120, 202. **(H)** Representative fluorescence images of cells containing nanobody coacervate hubs showing two topologies of GFP recruitment: surface or interior; higher in the +CQ case. Coacervates formed from 0.5mg/mL HBpep-SA co-assembled with 0.1mg/mL VHHGFP4-mCherry were added to the cell media with or without 100 µM Chloroquine. CQ refers to Chloroquine. Scale bar: 10µm. Source data are provided as a Source Data file.

To optimize the hub system and methodology we decided to generate a stable monoclonal human cell line expressing the GFP target and explored additional mediators of endosomal escape. Using the monoclonal cell line with more uniform expression we found reduced non-specific recruitment to the control hubs lacking nanobody; 0% hubs showing recruitment (Supplementary Fig. 5D) and 0% of cells with a recruiting hub (Fig. S5E). Whereas nanobody hubs still selectively bound to target: 13% of hubs showed targeting and 81% of cells had at least one recruiting hub. (Supplementary Fig. 5D, E). Second, we tested use of chloroquine (CQ), a molecule can partition to endosomes and act as a proton sponge that causes overuse of proton pumps and endosomal membrane rupture^44^. We added to the cell media during hub delivery and then imaged cells 1 day later. We found that addition of CQ had a number of useful effects related to target binding to hubs. First, it caused a large increase in the fraction of hubs that could recruit target protein to the surface or interior (Fig. 5G). Interestingly, most of the recruitment was to the surface, similar to our in vitro data in Fig 4. (Fig 5H). Targeting was still highly selective, as control hubs showed de-enrichment at the surface, whereas hubs co-assembled with VHHGFP4 nanobody showed target surface enrichment, with an enrichment index for GFP of up to 2-4 (Supplementary Fig. 5F-H). Combining surface and internal target binding to our hubs, we observed that 44.6% of hubs showed GFP targeting (Fig. 5G), a value that exceeds most reported values of LNP endosomal escape^42,43^. Nanobody loaded hubs are stable in cells over time, in terms of number and size, up to 5 days (Supplementary Fig. 5I, J). Interestingly, if waited up to 5 days after hub delivery with CQ, we saw that most of GFP recruitment to the hubs was now internal and was very efficient: up to 40% of hubs showed target enrichment (Supplementary Fig. 5K). These data show that with increased endosomal escape and time, hub targeting can be highly selectively and highly efficient.

Given that hubs delivered to cells could show both topologies of target recruitment – surface and internal – we wanted to re-examine target recruitment in vitro. In our in vitro experiments (Fig 4), VHHGFP4-mCherry loaded condensates were only able to recruit GFP to the surface, likely limited by the small coacervate pore size^39^. In cells, HBpep-SA condensates may partially disassemble inside acidifying endosomes or upon escape to the reducing environment of the cytosol, likely due to increased porosity, which would allow larger macromolecules such as GFP to diffuse in. To test this, we pre-assembled nanobody-coacervates in vitro and imaged GFP target recruitment to hubs following addition of glutathione (GSH) for reducing, or acetate pH 4.5. Both conditions promoted reconfiguration of target recruitment, from entirely at the hub surface, to now also inside the compartment (Supplementary Fig. 5L). Combined, these data show surface and internal recruitment are feasible both in vitro or in cells in vivo. Notably, the partial disassembly of HBpep-SA coacervates through GSH reduction is slow when GSH is at similar concentration to that in the cytosolic (10 mM), taking hours, further explaining the high stability and long duration of observation of HBpep-SA coacervates delivered to cells.

### Hubs as Functional Degradosomes from Co-Assembly with bioPROTACs

We next wanted to test the feasibility of hubs to serve as reaction centers to regulate intracellular targets. We sought to assemble degradation hubs, in which bioPROTACs were pre-loaded as cargoes inside HBpep-SA coacervates. bioPROTACs are fusion domains that are able to recruit target proteins and an E3 ligase complex in proximity to one another for ubiquitination and degradation^45^. However, bioPROTACs cannot readily cross the cell membrane and face challenges with efficiency when simply genetically expressed in the cytosol^46^. It has been suggested that multimerizing bioPROTACs by forming condensates can increase degradation efficiency^47^. Therefore, we reasoned that bioPROTACs could be more functional when housed within an HBpep microcompartment delivered to cells.

We selected a proven GFP bioPROTAC (VHHGFP4-SPOP_167-374_) for co-assembly in HBpep-SA coacervates and delivered them to our U2OS monoclonal GFP-expressing cell line to test the efficiency of target degradation (Fig. 6A)^48^. When co-assembled with the bioPROTAC construct (VHHGFP4-SPOP_167-374_), HBpep-SA coacervates efficiently recruited GFP into hubs and decreased cytosolic GFP levels by ~78% compared to control coacervates (Fig. 6B, C) after one day. Hubs containing only the GFP nanobody VHHGFP4 alone did not decrease the cytosol GFP level, indicating the decreased in cytosolic GFP signal in cells containing hub loaded bioPROTAC is due to GFP degradation and not simply GFP sequestration. When we loaded just the E3 adaptor portion (SPOP_167-374_) of the bioPROTAC, we observed modest GFP degradation, presumably by non-specific ubiquitination and degradation of targets. We noticed target enrichment to hubs still occurred when loaded with bioPROTAC (Fig 6B) even though overall cytosolic signal was greatly reduced, suggesting that ubiquitylated targets may be protected from degradation inside the hub. As noted above, we do not believe this is a partitioning effect alone as the nanobody loaded hubs did not show lowered cytosolic signal (Fig 6C). Rather we posit that when targets are ubiquitylated at hubs and dissociate back into the cytosol, they are then degraded by proteosome. Overall, these data shows that bioPROTAC-loaded HBpep-SA coacervates can function as ‘degradosomes’ that selectively target protein for degradation, offering a potential generalizable strategy for target and pathway modulation in cells.

**Figure 6.**
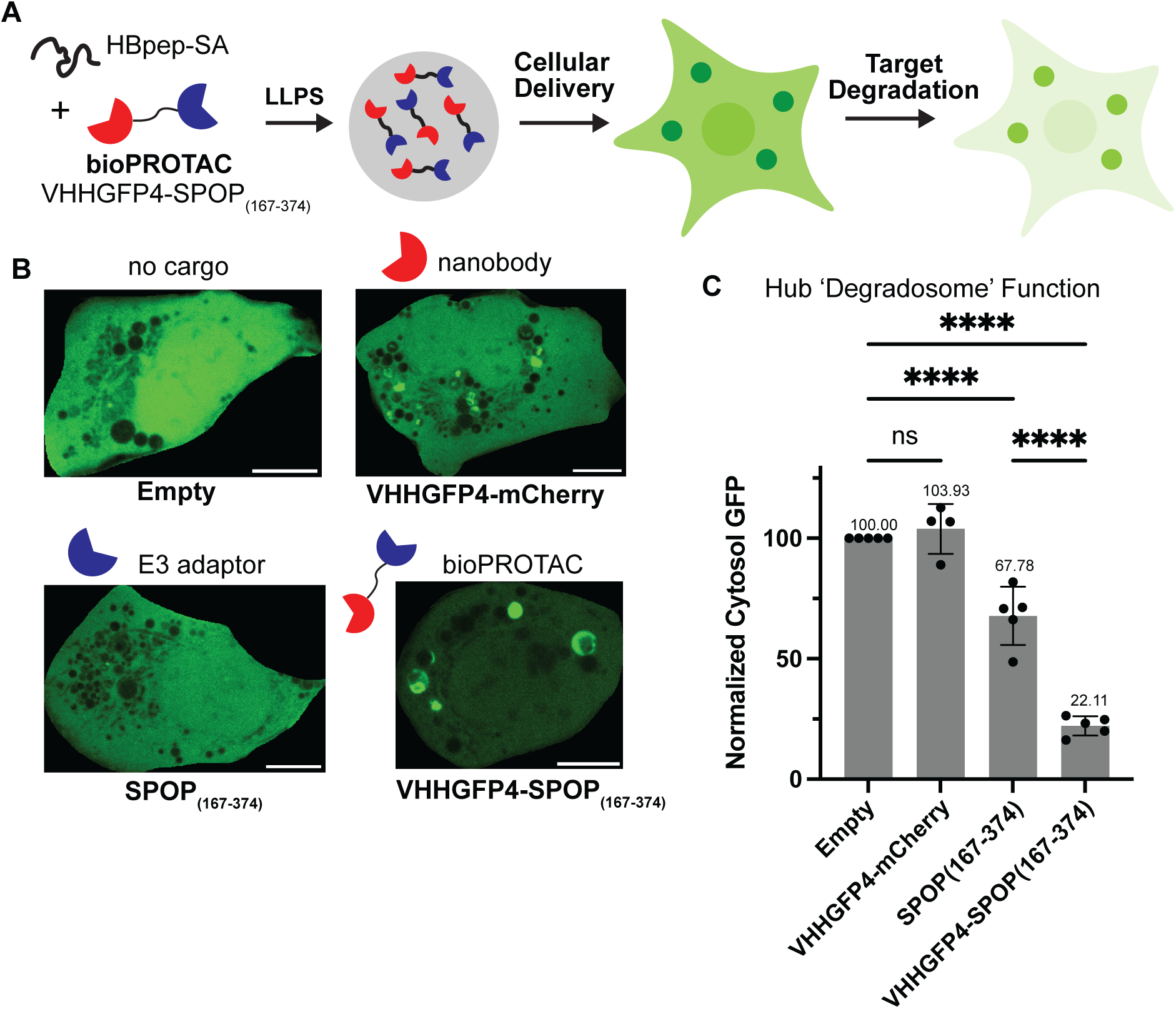
bioPROTAC-loaded hubs function as reaction centers for target degradation in cells. **(A)** Scheme showing bioPROTAC-loaded HBpep-SA coacervates delivered to GFP expressing cells and subsequent degradation of target GFP. **(B)** Representative fluorescence images set to same black and white values showing GFP flurecence in monoclonal GFP expressing cell line after delivery of coacervate hubs (preformed HBpep-SA coacervates containing either no cargo or 1.35 µM of the following: VHHGFP4 SPOP_167-374_ or bioPROTAC VHHGFP4-SPOP_167-374._ Scale bar: 10µm**. (C)** Quantification of normalized average cytosolic target GFP fluorescence intensity. n=5 repeats. Error bars indicate standard deviation from the mean. Center of error bars indicate the mean. Ordinary one-way ANOVA tests were used. Šidák correction is used to correct for multiple comparisons. P-value is 0.9251 for ns, and P ≤ 0.0001 for other comparisons. Source data are provided as a Source Data file.

We conclude it is feasible to deliver nanobody coacervates to cells that can function for days as stable hubs, capable of targeting endogenously expressed proteins to the microcompartment. These results demonstrate a one-pot solution toward remodeling protein interaction networks in living cells using synthetic hubs or organelles and represent a unique opportunity for cell engineering without the requirement for gene delivery.

## Methods

### Peptides and Proteins

HBpep and FITC-HBpep peptides were customize ordered from Genescript

HBpep-SA peptides were ordered from MCE (MedChemExpress, (HY-000)) and TargetMol (Catalog No. TP2797)

HBpep amino acid sequence^29^: GHGVY GHGVY GHGPY GHGPY GHGLY W

Tag-free GFP was purchased from abcam (ab84191, Recombinant A. victoria GFP protein)

### Oligonucleotides and Proteins

Oligonucleotide and protein sequences are listed in supplementary information.

### Cells

U2OS cells were acquired from ATCC as U-2 OS HTB-96™. B16 cells were gift from Andy Minn’s lab, which were purchased from ATCC as B16-F10 CRL-6475™. Human Primary Monocytes were purchased from Human Immunology Core at the Perelman School of Medicine at the University of Pennsylvania.

### Temperature shift method to induce LLPS

In this method, HBpep peptide powder was dissolved in 1x phosphate buffered saline (PBS) pH 7.4, aliquoted as a high concentration stock (~10mg/ml) and stored at −80C. To make coacervates, HBpep peptide and cargo were added to 1x PBS solution at 50°C and allowed to thoroughly mix, with all reagents in 0.5ml Eppendorf tube in a bead bath. Following 10 mins of incubation, the tube was shifted to room temperature (~23°C) for 15 mins to induce formation of coacervates.

### pH shift method to induce LLPS

In this method, HBpep and HBpep-SA peptide powder were dissolved in 50 mM acetate buffer at pH 4.5, aliquoted as high concentration stocks(~15mg/ml) and stored at −80°C. To generate coacervates, dissolved HBpep or HBpep-SA peptides had added to them sequentially water, 1M NaCl salt and cargo, and after thorough mixing, 10x phosphate buffer was added to shift the pH to ~7.4. We waited 15 minutes at room temperature for coacervate particles to formation and gelate, following the pH shift. The final solution is expected to have 100ml NaCl, 10.1mM Na_2_HPO_4_, 1.8mM KH_2_PO_4_ and 5mM Acetate besides peptide and cargo.

### Delivery of condensates to cells

Cells were seeded at 50% confluency one day before delivery in glass bottom 96-well plates (Cellvis P96-1-N). At this time, a final concentration of 10µg/mL DAPI was added in the culture to stain nucleus. On the day of coacervate delivery, cells were first washed with growth medium: EMEM for U2OS cells, DMEM for B16-F10 melanoma cells and RPMI1640 for primary human monocytes. For coacervate delivery, growth medium was replaced by a 100µl solution of Opti-MEM combined with a volume of 10-30 µl of preformed coacervates. Particle uptake was detectable within 4 hr and cells. We generally quantified at 24 hr after coacervate addition, following extensive washing of the wells to remove and external particles. Membranes were stained by Abcam Cytopainter Cell Membrane Staining Kit in the experiment in Figure 3A. For stability tests in Figure 3B-E, Figure S5I,J, U2OS cells were cultured in EMEM for 5 days. For Figure5G,H, Chloroquine was added in growth medium at 100µM final concentration. For Figure S5K, Chloroquine was added in the growth medium for 5 days at 20µM final concentration.

### Partial disassembly of HBpep-SA coacervates in vitro by Acid and GSH in vitro

To test feasibility of partially disassembling HBpep-SA coacervates by acidification and redox state we tested titration to imaging wells of acetate pH 4.5 and GSH. We first added 30µl of preformed HBpep-SA coacervates (assembled at 0.5 mg/mL) loaded with 0.1mg/mL VHHGFP4-mCherry on top of 100 µl Opti-MEM media in glass-bottom 96 well-plates. After waiting 15 mins and confirming visible formation of coacervates via microscopy, we then titrated the solution by adding 500 mM Acetate pH4.5 in 1µl volume increments (pipetted on top of the well solution). Generally, partial disassembly of HBpep-SA coacervates – detectable by internal recruitment of GFP target - was observed after ~1 or 2µl titration and within minutes. For testing the effect of GSH on partial disassemble HBpep-SA coacervates, we performed a similar set of steps. We added 30µl of preformed HBpep-SA coacervates (assembled at 0.5 mg/mL) loaded with 0.1mg/mL VHHGFP4-mCherry on top of 100µl Opti-MEM in glass-bottom 96 well-plates. After waiting 15 minutes, we added GSH to a final concentration at 10 mM and incubated the plates at 37°C for ~2.5 hours. Partial disassembled HBpep-SA coacervates – detectable by internal recruitment of GFP target – were observed primarily among smaller coacervates.

### Cloning

Codon-optimized 6his-GFPHBP, 6his-mCherry, 6his-VHHGFP4-mCherry, 6his-VHHGFP4-mCherry-HBP, 6his-VHHGFP4-SPOP_167-374_ and 6his-SPOP_167-374_ constructs were cloned into pET expression vectors between NdeI and XhoI sites and assembled via In-Fusion ligation (Takara Bio) for transformation into Stellar competent cells. For stable cell lines, GFP was cloned into pLJM1 vector between AgeI and EcoRI sites and assembled in Stbl3 competent cells by In-Fusion ligation (Takara Bio).

### Protein purification

In all cases, recombinant His-tagged proteins were purified from E. coli lysates using Ni-NTA agarose beads and affinity chromatography. First, pET plasmids were transformed into Rosetta 2 Escherichia coli cells (Sigma Aldrich). Bacterial cultures were grown in Luria Broth (LB) supplemented with kanamycin and chloramphenicol at 37 °C to an OD600 of 0.6–0.8, and expression was induced by addition of 0.5 mM isopropyl β-D-1-thiogalactopyranoside. Cultured cells were grown overnight a 16 °C overnight (18hr). Cell cultures were pelleted by centrifugation at 1,960g and the resulting pellets were stored at −80 °C. To generate cell lysates, bacterial pellets were thawed, then resuspended in lysis buffer (50 mM Tris-HCl, pH 7.5, 1 M NaCl, 20 mM imidazole, 1 mM β-mercaptoethanol) along with complete EDTA-free protease inhibitor cocktail (Roche). Cells were lysed with 2 minutes of sonication at 50% power using a Branson Sonifier. Lysates were clarified by centrifugation at 13,250g for 20 min in a Sorvall RC6+ centrifuge using a F21S-8x50y rotor (Thermo Fisher Scientific) at 4 °C and the supernatant was collected. For soluble proteins, the supernatant was incubated with Ni–NTA beads (Thermo Fisher Scientific) while rotating at 4°C for 1 h. Beads were then washed three times. Proteins were eluted with elution buffer (20 mM Tris-HCl, pH 7.5, 500 mM imidazole, 1 mM dithiothreitol (DTT), and 1M NaCl). Eluted proteins were dialyzed overnight into PBS buffer (10mM Na2HPO4,1.8mM KH2PO4, 2.7mM KCl, 137 mM NaCl, pH7.4) using a Slide-A-Lyzer membrane cassette (Thermo Fisher Scientific) with a 10-kDa size cutoff at room temperature^49^. Protein aliquots were flash frozen and stored at −80°C.

For 6his-VHHGFP4-SPOP_167-374_, which was insoluble, the pellet after clarification was dissolved in urea, the protein purified via affinity on Ni-NTA beads and then slowly refolded. Pellets were re-suspended in buffer (6M Urea, 50mM Tris-HCl, pH7.5, 1M NaCl, 20mM imidazole) and this re-suspension solution was centrifuged at 13250g for 20min. The collected supernatant was then incubated with Ni-NTA beads while rotating at 4 °C for 1 h. Beads were then washed three times. Proteins were eluted by elution buffer (20mM Tris-HCl, pH 7.5, 500 mM imidazole, 1 mM dithiothreitol (DTT), and 1M NaCl, 6M Urea). The elution solution was then dialyzed for a total of 3 days with stready graded reducection urea concentration from 6 M to 4 M, 2 M, 1 M, 0.5 M and 0M (in PBS buffer) and finally dialyzed into PBS buffer.

### Mammalian cell procedures

U2OS cells were cultured in Eagle’s minimal essential medium (EMEM, Quality Biological) supplemented with 10% fetal bovine serum (Gibco), 2 mM L-glutamine (Gibco) and 10 U mL−1 penicillin-streptomycin (Gibco). B16 cells were cultured in Dulbecco’s Modified Eagle’s Medium supplemented with 10% fetal bovine serum (Gibco), 2 mM L-glutamine (Gibco) and 10 U mL−1 penicillin-streptomycin (Gibco). Cells are maintained at 37 °C in a humidified atmosphere with 5% CO2. Cells were split in 1:5 ratio every three days and had been passaged for less than two months, were negative for known infection. Frozen human primary monocytes were obtained from Penn Human Immunology Core, then thawed and cultured in RPMI1640 with 10% fetal bovine serum (Gibco), 2 mM L-glutamine (Gibco) and 10 U mL−1 penicillin-streptomycin (Gibco).

### Generating stable cell lines

At day 0, wildtype U2OS cells were seeded in 24-well plates at 70-90% confluence in 500µl Eagle’s minimal essential medium (EMEM, Quality Biological) supplemented with 10% fetal bovine serum (Gibco), 2 mM L-glutamine (Gibco) and 10 U mL−1 penicillin-streptomycin (Gibco). At day 1, growth medium was replaced with 500 µl GFP lentivirus stock. At day 2, growth medium was replaced back to EMEM with 10% FBS, 2mM L-glutamine and 10 U mL−1 penicillin-streptomycin. At Day 4, cells were trypsinized and collected for Fluorescence-activated cell sorting (FACS) to select GFP-expression cells in the middle of the expression distribution for generating polyclonal U2OS cell line. This polyclonal cell line was expanded for 2 weeks and stored in EMEM with 10% DMSO in nitrogen tank.

To generate monoclonal GFP-expressing U2OS cell line, the polyclonal cell line in culture was trypsinized and collected for Fluorescence-activated cell sorting (FACS) for medium GFP expression cells. 96 single U2OS cells were sorted in 96 well-plates (one cell per well). After 2 weeks of expansion, 4 mono-clonal cell lines were generated and stored in EMEM with 10% DMSO in nitrogen tank. A monoclonal line expressing low-medium GFP levels was chosen for experiments in Figure 5H, S5C-K and Figure 6.

### Microscopy

Images were collected on an Olympus IX81 inverted confocal microscope (Olympus Life Science) equipped with a Yokogawa CSU-X1 spinning disk, a mercury lamp, 405- 488- and 561-nm laser launches, and an iXon3 electron-multiplying charge-coupled device (EMCCD) camera (Andor). Multidimensional acquisition was controlled by MetaMorph software (Molecular Devices, version 7.8.10.0). Samples were illuminated using a 405-nm laser and/or 488-nm laser and/or a 561-nm laser and were imaged through a ×100, 1.4-NA oil immersion objective and/or a x20, 0.75NA air objective. Z stacks were collected in 0.3-μm steps.

For in vitro experiments, HBpep and HBpep-SA condensates were imaged in custom-fabricated acrylic gasket chambers adhered to glass coverslips (#1.5 glass thickness; Corning Inc.) The glass bottom was passivated by 10mg/mL BSA overnight. For in vivo experiments, cells were seeded in glass bottom 96-well plates (Cellvis P96-1-N).

For FRAP experiments on FITC-HBpep labeled HBpep and HBpep-SA condensates, a Roper iLas2 photoactivation system controlling a 405-nm laser was used. An area containing multiple condensates were selected and photobleached, and fluorescence recovery in the bleached region was analyzed in ImageJ after normalizing the fluorescence intensity from 0 to 100 based on the florescence intensity before FRAP and right after FRAP. Images were collected to visualize the condensates at the 488-nm wavelength and bright field using a ×100, 1.4-NA oil immersion objective.

### Image analysis

Image segmentation and analysis were performed in FIJI and Excel. “Analyze Particle” function in FIJI was used to count and measure the size of condensates in Figure 1 and Figure S1 and measure the average fluorescence intensity in condensates in Figure 2 (except for temperature based assembly with GFP). In other quantitation, measures of condensate size, number and fluoresce intensity were measured manually in FIJI. Calculation of Enrichment Index in Figure 2 for cargo loading of GFP and GFPHBP was calculated by measuring mean fluorescence intensity of whole coacervate divided by average fluorescence intensity in the continuous phase outside coacervate. EI= (Intensity_coacervate_/Intensity_solution_). For line scans, such as in Figure 4, these were performed using FIJI in the 488 nm channel. Maximum fluorescence intensity was recorded and divided by the average fluorescence intensity in the liquid phase outside condensates to determine ring enrichment index. To measure target partitioning and enrichment to hubs In Figure 5E-F and Figure S5A-B, line scans were performed in FIJI in the 488 nm channel. The maximum fluorescence intensity was recorded and divided by the average fluorescence intensity in the cell cytosol to generate the maximum enrichment index. EI_max_= (Max.Intensity_coacervate_/Average.Intensity_cytosol_). Similarly for Figure 5G, Figure S5C, D, the mean enrichment index was calculated by dividing average coacervate GFP fluorescence intensity was divided by the mean fluorescence intensity recorded in FIJI. For target surface enrichment calculations in In Figure 5G, Figure S5G, the average surface GFP fluorescence intensity derived from a line scan was divided by the mean cytosol GFP fluorescence intensity using FIJI. The 3D reconstruction in the Supplement Movie2 was rendered in Imaris. For measurements of degradation activity in Figure 6C, the average cytosol GFP fluorescence intensity was measured in FIJI by selecting multiple random cytosolic areas excluding hubs and and nucleus.

### Statistics and reproducibility

The experiments were reproducible. All statistical analyses were performed in GraphPad Prism 9. To test the significance of two categories, an unpaired one-tailed Welch’s t-test was used. To test the significance of more than two categories, a one-way Brown-Forsythe ANOVA was used. In all cases, NS indicates not significant (P > 0.05), *P ≤ 0.05, **P ≤ 0.01, ***P ≤ 0.001 and ****P ≤ 0.0001. The sample size was determined by sample and image acquisition. Sample images were acquired randomly within an imaging well. Cells and coacervates at the edges of images were excluded. The Investigators were not blinded during experiments and outcome assessment.

## Data Availability

The data supporting the findings of this study are available within the Article and Supplementary Information. Raw images are available from the corresponding author upon reasonable request. Source data are provided as a Source Data file.

## Acknowledgements

We thank A. Stout and the Penn CDB Microscopy Core for imaging support, and M. Wadley for human primary monocytes cell culture, and W. Wentao for support of GFP cell line generation. We thank the Human Immunology Core and the Division of Transfusion Medicine and Therapeutic Pathology at the Perelman School of Medicine at the University of Pennsylvania for providing de-identified, primary human monocytes. This study was partially supported by the National Institute of Biomedical Imaging and Bioengineering grant, EB028320 (M.C.G.), by National Science Foundation through the University of Pennsylvania Materials Research Science and Engineering Center (MRSEC) (DMR-2309043), and a seed award from the Center for Precision Engineering for Health (CPE4H) at the University of Pennsylvania (M.C.G., A.J.P.)

## Author Contributions

M.C.G, W.T. and A.J.P. conceptualized the project. M.C.G., W.T. R.Q.T., D.M.C. designed the experiments. W.T. and P.J. performed initial peptide synthesis. W.T. and R.Q.T. designed and cloned genetic constructs. W.T. purified cargo and nanobody proteins and performed experiments for assembly, co-assembly, FRAP, in vitro targeting experiments and coacervate stability experiments. W.T. and R.Q.T performed imaging for characterization of coacervate distribution and mobility in various cell types. W.T. performed experiments for GFP targeting in cells. W.T. analyzed experimental data. M.C.G and W.T. wrote the manuscript with feedback from A.J.P. and D.M.C.

## Competing Interests

The authors declare no competing interests.

## Materials and Correspondence

Plasmids will be made available upon request to Prof. Matthew Good

## Notes

### Competing Interest Statement

The authors have declared no competing interest.

### Summary of Updates

Added new results in the form of figure panels 1. New Figure 6 - bioPROTAC loaded hubs efficiently degrade target proteins in cells 2. Updates with new experiments in: Figure 4, Figure 5. Also in Supplemental FIg S1, S2, S4, S5. Revised text to accompany additional new experimental results.

