## Supplementary Materials PDF for "Delivery of Peptide Coacervates to Form Stable Interaction Hubs in Cells"

**Supporting Information Includes:**

List of Proteins Used: Page 1
List of Oligonucleotides Used: Page 2
Supplementary Movie Legends: Pages 3
Supplementary Figures S1-S5: Pages 4-10

**List of Proteins Sequences Used**

| Protein Name | Protein Sequence |
|---|---|
| 6His-TEV-EGFP-HBP | MHHHHHHENLYFQSMVSKGEELFTGVVPILVELDGDVNGHKFSVSGEG EGDATYGKLTLKFICTTGKLPVPWPTLVTTLTYGVQCFSRYPDHMKQHDF FKSAMPEGYVQERTIFFKDDGNYKTRAEVKFEGDTLVNRIELKGIDFKED GNILGHKLEYNYNSHNVYIMADKQKNGIKVNFKIRHNIEDGSVQLADHYQ QNTPIGDGPVLLPDNHYLSTQSALSKDPNEKRDHMVLLEFVTAAGITLGM DELYKGHGVYGHGVYGHGPYGHGPYGHGLY |
| mCherry-6His | MVSKGEEDNMAIIKEFMRFKVHMEGSVNGHEFEIEGEGEGRPYEGTQT AKLKVTKGGPLPFAWDILSPQFMYGSKAYVKHPADIPDYLKLSFPEGFKW ERVMNFEDGGVVTVTQDSSLQDGEFIYKVKLRGTNFPSDGPVMQKKTM GWEASSERMYPEDGALKGEIKQRLKLKDGGHYDAEVKTTYKAKKPVQL PGAYNVNIKLDITSHNEDYTIVEQYERAEGRHSTGGMDELYKHHHHHH |
| 6His-VHHGFP4-mCherry | MHHHHHHMVQLVESGGALVQPGGSLRLSCAASGFPVNRYSMRWYRQA PGKEREWVAGMSSAGDRSSYEDSVKGRFTISRDDARNTVYLQMNSLKP EDTAVYYCNVNVGFEYWGQGTQVTVSSVSKGEEDNMAIIKEFMRFKVH MEGSVNGHEFEIEGEGEGRPYEGTQTAKLKVTKGGPLPFAWDILSPQFM YGSKAYVKHPADIPDYLKLSFPEGFKWERVMNFEDGGVVTVTQDSSLQ DGEFIYKVKLRGTNFPSDGPVMQKKTMGWEASSERMYPEDGALKGEIK QRLKLKDGGHYDAEVKTTYKAKKPVQLPGAYNVNIKLDITSHNEDYTIVE QYERAEGRHSTGGMDELYK |
| 6His-VHHGFP4-mCherry-HBP | MHHHHHHMVQLVESGGALVQPGGSLRLSCAASGFPVNRYSMRWYRQA PGKEREWVAGMSSAGDRSSYEDSVKGRFTISRDDARNTVYLQMNSLKP EDTAVYYCNVNVGFEYWGQGTQVTVSSVSKGEEDNMAIIKEFMRFKVH MEGSVNGHEFEIEGEGEGRPYEGTQTAKLKVTKGGPLPFAWDILSPQFM YGSKAYVKHPADIPDYLKLSFPEGFKWERVMNFEDGGVVTVTQDSSLQ DGEFIYKVKLRGTNFPSDGPVMQKKTMGWEASSERMYPEDGALKGEIK QRLKLKDGGHYDAEVKTTYKAKKPVQLPGAYNVNIKLDITSHNEDYTIVE QYERAEGRHSTGGMDELYKGHGVYGHGVYGHGPYGHGPYGHGLY |
| 6His-VHHGFP4-SPOP$_{167-374}$ | MHHHHHHVQLVESGGALVQPGGSLRLSCAASGFPVNRYSMRWYRQAP GKEREWVAGMSSAGDRSSYEDSVKGRFTISRDDARNTVYLQMNSLKPE DTAVYYCNVNVGFEYWGQGTQVTVSSSGSVNISGQNTMNMVKVPECR LADELGGLWENSRFTDCCLCVAGQEFQAHKAILAARSPVFSAMFEHEME ESKKNRVEINDVEPEVFKEMMCFIYTGKAPNLDKMADDLLAAADKYALE RLKVMCEDALCSNLSVENAAEILILADLHSADQLKTQAVDFINYHASDVLE TSGWKSMVVSHPHLVAEAYRSLASAQCPFLGPPRKRLKQS |
| 6His-SPOP$_{167-374}$ | MHHHHHHVNISGQNTMNMVKVPECRLADELGGLWENSRFTDCCLCVA GQEFQAHKAILAARSPVFSAMFEHEMEESKKNRVEINDVEPEVFKEMMC FIYTGKAPNLDKMADDLLAAADKYALERLKVMCEDALCSNLSVENAAEILI LADLHSADQLKTQAVDFINYHASDVLETSGWKSMVVSHPHLVAEAYRSL ASAQCPFLGPPRKRLKQS |

**List of Oligonucleotides Used**

| Primer Name | Primer Sequence |
|---|---|
| 6His F | 5’-AAGGAGATATACATATGCACCACCACCACCA-3’ |
| HBpep-stop R | 5’-GGTGGTGGTGCTCGAGTTATTAATATAAGCCGTGCCCGTAC-3’ |
| mCherry R | 5’-GGTGGTGGTGCTCGAGTTAtTACTTGTACAGCTCGTCCATGC-3’ |
| SPOP167-374 R | 5’-GGTGGTGGTGCTCGAGTTAtTAAGATTGTTTCAGACGTTTGCGAG-3’ |

**Supplementary Movie Legends**

**Supplementary Movie M1: Cytoplasmic coacervates show Brownian motion in cells.** Time lapse of HBpep-SA condensates moving inside U2OS cells. 10µl of 2 mg/mL HBpep-SA condensates labeled by FITC were delivered to U2OS cells and cells were imaged 28 hrs after delivery. The Movie stands for an actual duration of 40 seconds timelapse with 2 seconds interval per frame. Scale bar: 10µm.

**Supplementary Movie M2: Cell 3D reconstruction shows uptake of many large peptide coacervates.** 3D reconstruction of a B16-F10 cell filled with HBpep-SA condensates loaded with GFP-HBP. Titling of 3D reconstruction performed in Imaris from an image stack.

### Supplementary Figures

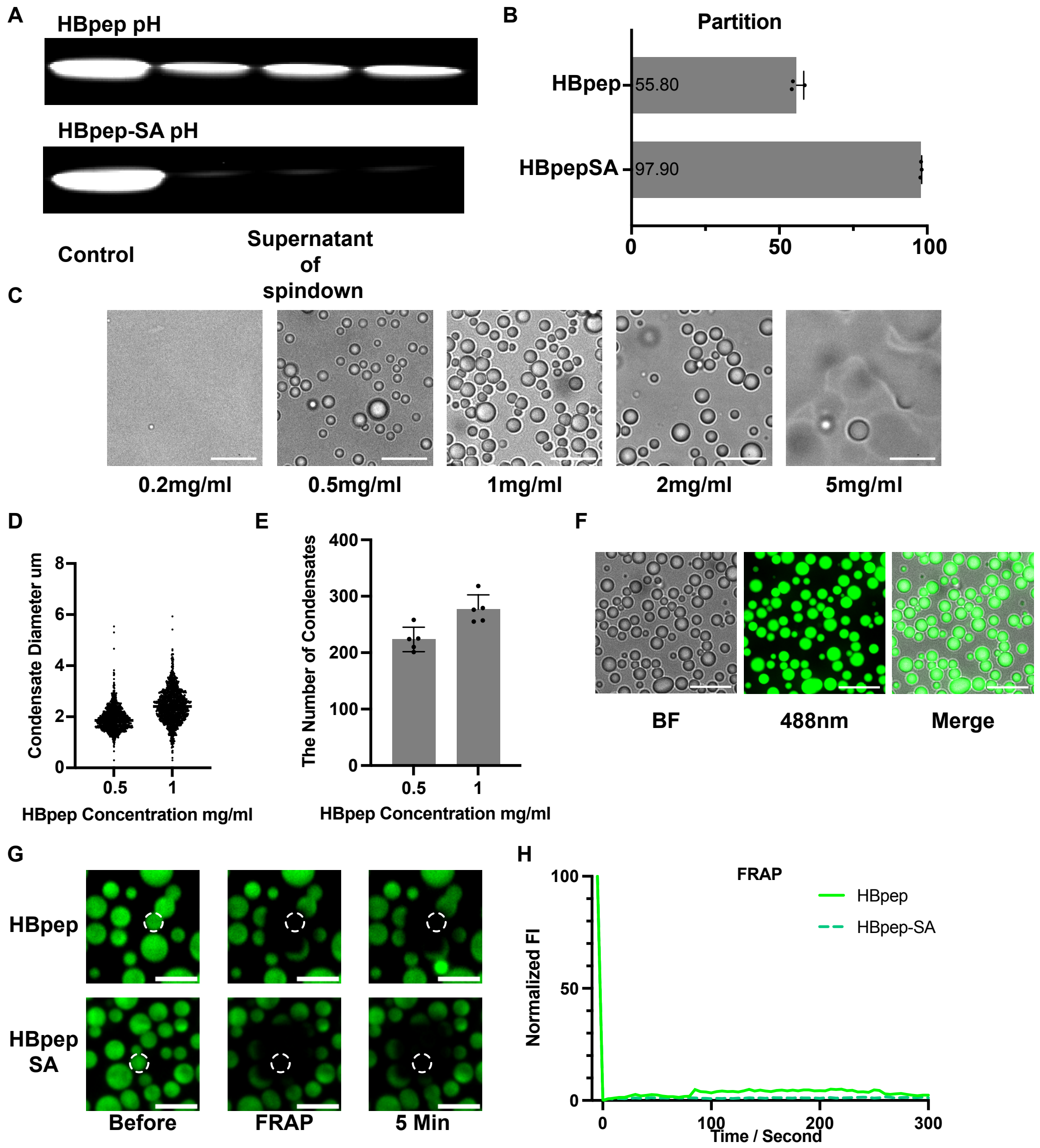


**Supplementary Figure S1: Further characterization of size control for HBpep and HBpep-SA coacervates assembled via temperature shift.**

**(A)** SDS-PAGE gel showing protein remaining in supernatant following sedimentation of coacervate formulations. From assembly of 0.5mg/mL HBpep and HBpep-SA using pH shift method. **(B)** Quantification of HBpep and HBpep-SA partitioning to coacervate phase from sedimentation analysis via SDS-PAGE gel; from S1A. Error bars indicate standard deviation. Center of error bars indicate means. **(C)** Brightfield microscopy images showing assembly of HBpep coacervates at different peptide concentrations using the temperature shift method: solution contains 100mM NaCl and phosphate buffer pH 7.4. Scale bar: 10 μm. **(D)** Quantification of HBpep condensate size (diameter) when assembling at different peptides concentrations via temperature shift method**.** n=1118, 1385, **(E)** Quantification of HBpep condensate number per imaging area: 4616.2 $\mu m^2$. n=5. Error bars indicate standard deviation. Center of error bars indicate means. **(F)** Fluorescence images of FITC-HBpep labeled HBpep condensates (1:100) assembled at 1 mg/mL peptide concentration using temperature shift. scale bar: 10 μm. **(G)** Fluorescence images over time following laser photobleaching, showing FITC labeled (1:100) HBpep and HBpep-SA coacervates formed at 0.5 mg/mL via pH shift, Scale bar: 5 μm. **(H)** Quantification of normalized fluorescence recovery after photobleaching showing intensity within bleached region of FITC labeled HBpep and HBpep-SA coacervates formed via pH shift. Source data are provided as a Source Data file.

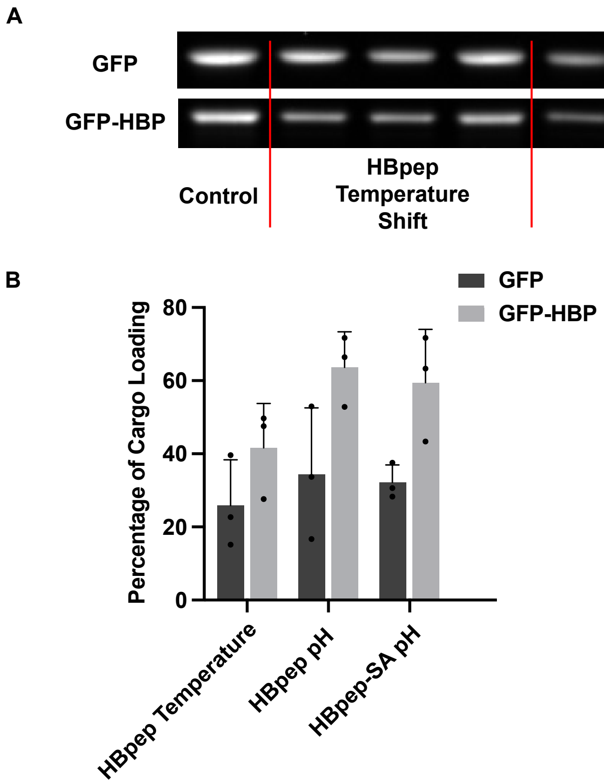


**Supplementary Figure S2: Quantitation of cargo loading within coacervates using particle sedimentation analysis.**

**(A)** SDS-PAGE gel showing cargo (GFP or GFP-HBP) remaining in the supernatant following spin down of coacervates. 0.1 mg/mL GFP or GFPHBP were co-assembled with 0.5mg/mL HBpep or HBpep-SA through either temperature shift or pH shift methods. Top gel: GFP; lower gel: GFP-HBP. Compared to loading control (left). **(B)** Quantification of cargo loading into coacervates based on SDS-PAGE gel analysis in S2A. (100% - percent remaining in supernatant). n=3. Error bars indicate standard deviation. Center of error bars indicate means. Source data are provided as a Source Data file.

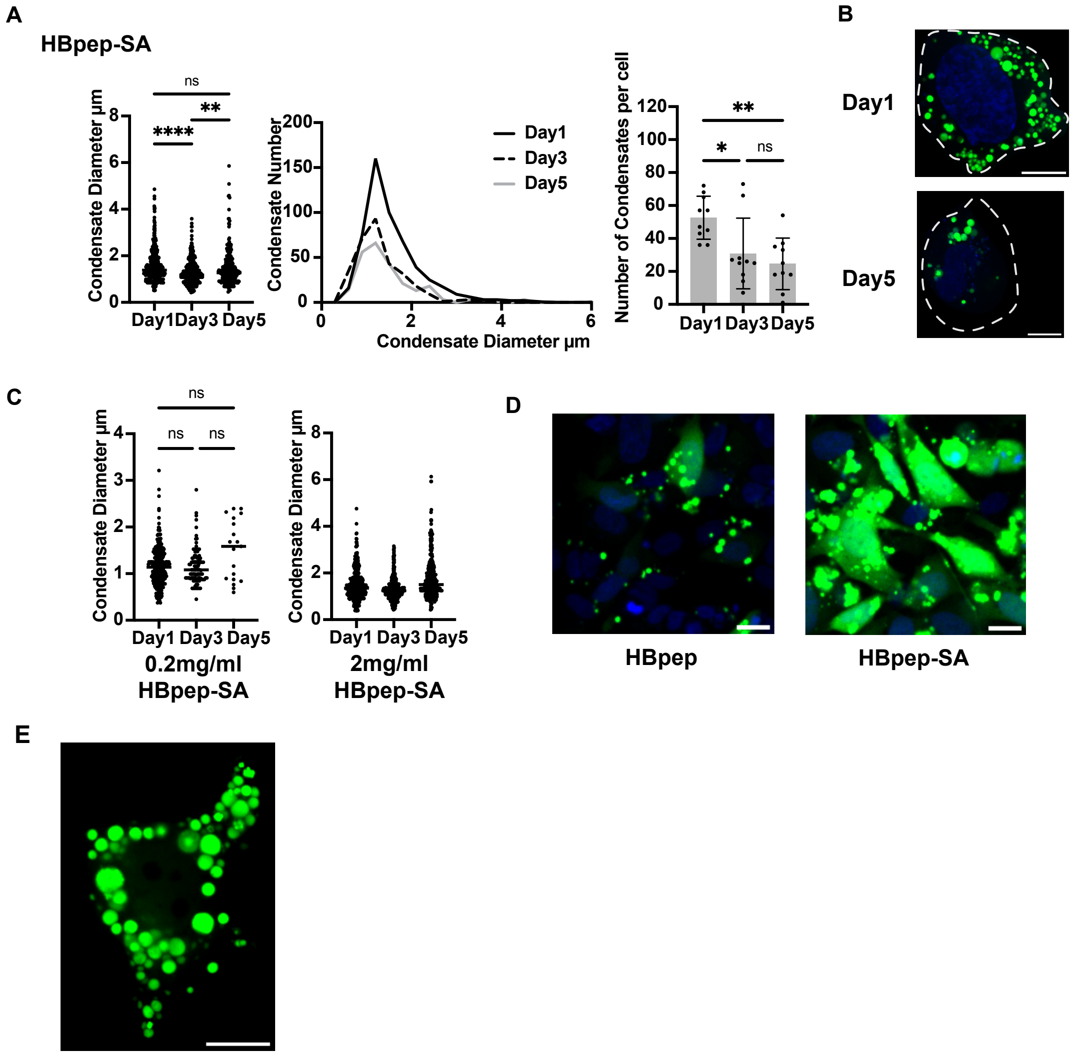


**Supplementary Figure S3: Efficiency of coacervate delivery to cells and stability over time.**
**(A)** Quantification of coacervate stability within U2OS cells over 5 days. Size and number of coacervates. Cells were initiated supplemented with particles from a formulation of 30 µl of 0.5mg/mL HBpep-SA, labeled by FITC-HBpep (1:100). Condensate number, n=526, 309, 246. Cell number, n=10. Brown-Forsythe and Welch One-Way ANOVA tests were used. Dunnett T3 statistical hypothesis is used to correct for multiple comparisons. Adjusted p-values are (for left) <0.0001, 0.0030, 0.5089 and (for right) 0.0443, 0.0013, 0.8365. Error bars indicate standard deviation. Center of error bars indicate means. **(B)** Representative fluorescence images of U2OS cells containing HBpep-SA coacervates on day 1 and day 5. Green: FITC labeled HBpep-SA condensates. Blue: Nucleus, stained DNA (DAPI). Scale bar: 10 µm. **(C)** Quantification of coacervate size within cells, comparing particle formulations at 0.2 mg/mL or 2mg/mL HBpep-SA. Coacervate number n =255, 108, 19(left), 364, 400, 258(right). Brown-Forsythe and Welch One-Way ANOVA tests were used. Dunnett T3 statistical hypothesis is used to correct for multiple comparisons. Adjusted p-values are 0.9997, 0.0828, 0.0822. **(D)** Representative fluorescence images of U2OS cells 24 hrs after being supplemented with 30 µl of HBpep or HBpep-SA coacervates assembled at 0.5mg/mL and loaded with 0.1mg/mL GFP-HBP. HBpep-SA coacervates release more cargo to the cytosol than HBpep coacervates. Green: GFP-HBP. Blue: Nucleus, stained DNA (DAPI). Scale bar: 20 µm. **(E)** Partial z-projection of an exemplary cell that has taken up approximately 100 condensates containing GFP-HBP. Scale bar: 10 µm. A portion of cell can be seen rotated in 3D in Supp Movie M2. NS indicates not significant (P > 0.05), *P ≤ 0.05, **P ≤ 0.01, ***P ≤ 0.001 and ****P ≤ 0.0001. Source data are provided as a Source Data file.

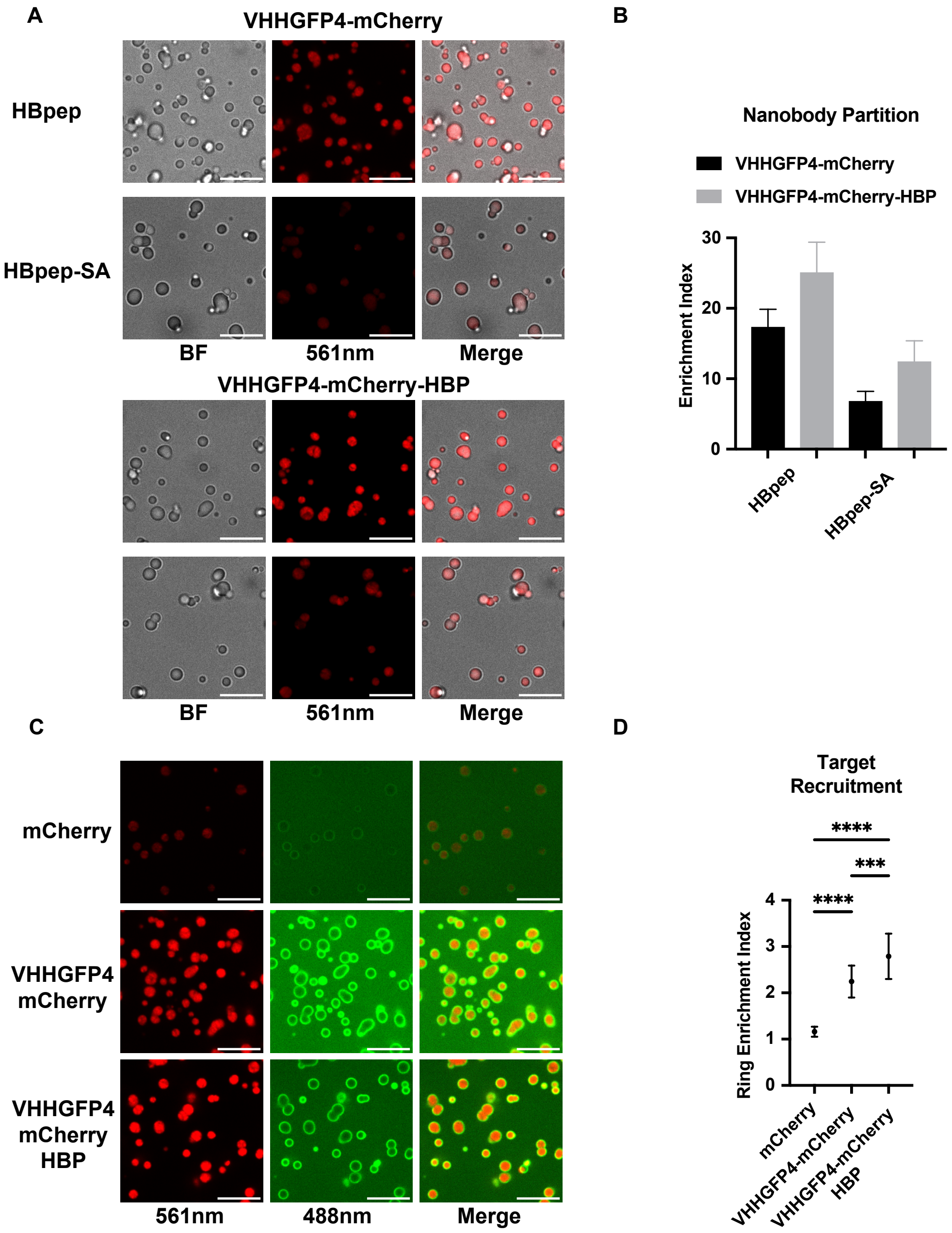


**Supplementary Figure S4: Further characterization of targeting function of nanobody-containing coacervates.** **(A)** Bright field and fluorescence images of nanobody loading in coacervates co-assembled from 0.5mg/mL HBpep and HBpep-SA with 0.1mg/mL VHHGFP4-mCherry or VHHGFP4-mCherry-HBP nanobodies. Scale Bar: 10 μm. **(B)** Quantification of nanobody loading in HBpep and HBpep-SA coacervates. Enrichment index: fluorescent intensity in coacervate divide by continuous phase. n=144, 90, 51, 65. Center of error bars indicate means. **(C)** Fluorescence images showing target binding to control (mCherry) versus nanobody containing coacervates. 1 μM final concentration of GFP was added to an imaging well containing HBpep condensates. scale bar: 10μm. **(D)** Quantification of the target (GFP) enrichment and partitioning to control versus nanobody containing HBpep coacervates. Measuring surface fluorescence intensity enrichment relative to the continuous phase. n=20. Brown-Forsythe and Welch One-Way ANOVA tests were used. Dunnett T3 statistical hypothesis is used to correct for multiple comparisons. Adjusted p-values are $<0.0001$, 0.0008, $<0.0001$. NS indicates not significant ($P > 0.05$), *$P \leq 0.05$, **$P \leq 0.01$, ***$P \leq 0.001$ and ****$P \leq 0.0001$. Error bars indicate standard deviation. Center of error bars indicate means. Source data are provided as a Source Data file.

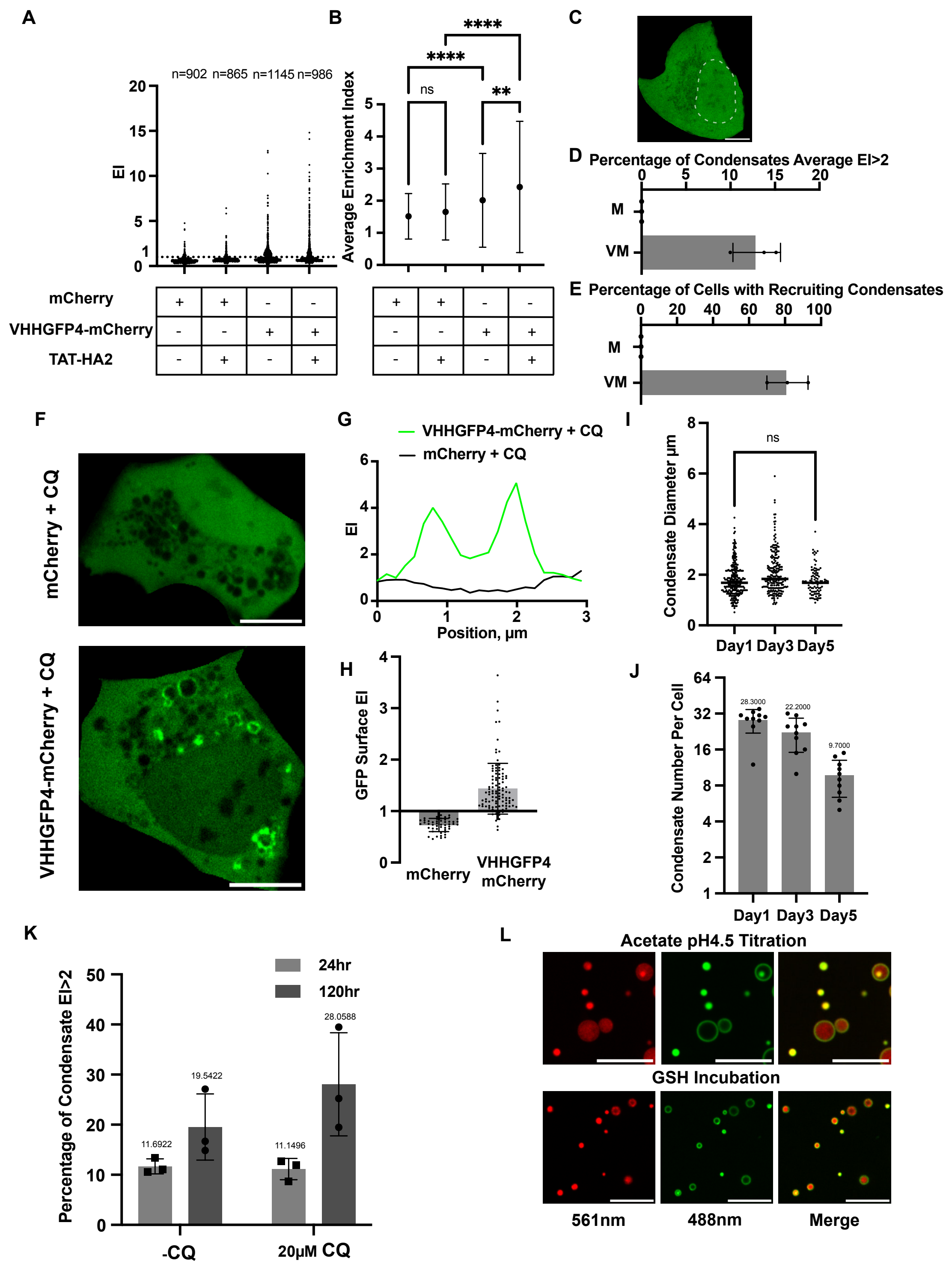


**Supplementary FigureS5. Distribution of 'hub' sizes following cellular uptake of coacervates, in presence an absence of assist peptide.**

**(A)** Quantification of the maximum target GFP enrichment in individual synthetic hubs within cells. **(B)** Quantification of the average maximum enrichment index (EI) – hub/cytosol signal - of target GFP inside hubs in cells, limited to those than show any partition coefficient above 1. n= 67, 86, 394, 315. Brown-Forsythe and Welch One-Way ANOVA tests were used. Dunnett T3 statistical hypothesis is used to correct for multiple comparisons. Adjusted p-values are <0.0001, 0.7473, 0.0097, <0.0001. NS indicates not significant ($P > 0.05$), $^*P \leq 0.05$, $^{**}P \leq 0.01$, $^{***}P \leq 0.001$ and $^{****}P \leq 0.0001$. Center of error bars indicate means. **(C)** A representative image of monoclonal U2OS cell line expressing genomeencoded GFP showing uniform-distributed fluorescence of GFP protein in cytosol. Scale bar: 10μm. **(D)** Quantification of target recruitment as percentage of delivered HBpepSA coacervates that show hub GFP enrichment index (EI) > 2 within monoclonal GFP U2OS cell line. Comparing control mCherry (M) loaded vs. nanobody VHH4-mCherry (VM) loaded hubs. n=3. (**E**) Quantitation of percentage of cells loaded with HBpepSA hubs that have at least one condensate with average EI > 2 for the monoclonal GFP U2OS cell line. n=3 repeats. Comparing control mCherry (M) loaded vs. nanobody VHH4-mCherry (VM) loaded hub. **(F)** Representative images showing hub surface recruitment of target to control vs. nanobody loaded hubs in presence of CQ. Scale bar: 10μm. (**G**) Line scan of GFP target enrichment to hubs in GFP expressing cells loaded with hubs containing mCherry or VHHGFP4-mCherry in the presence of CQ**. (H)** Quantification of surface enrichment index of condensates loaded with mCherry or VHHGFP4-mCherry. n=50, and n=107. **(I)** Quantification of hub size distribution in U2OS cells for HBpep-SA coacervates (30μl volume added to media) loaded with 0.1mg/ml VHHGFP4-mCherry after 1, 3 and 5 days. n=283, n=222, n=96. **(J)** Quantification of the number of hubs per U2OS cell for HBpep-SA coacervates (30μl volume added to media) loaded with 0.1mg/ml VHHGFP4-mCherry after 1, 3 and 5 days. n=10. **(K)** Quantification of percentage of HBpepSA hubs showing target recruitment with average EI > 2, comparing 1 and 5 days of delivery and +/- CQ. 30μl HBpep-SA loaded with 0.1mg/ml VHHGFP4-mCherry were delivered to polyclonal GFP-expressing U2OS cells. Note, a more modest concentration, 20μM, of CQ was added in the media to reduce cell cytotoxicity across 5 days. n=3 repeats **(L)** Representative images in vitro showing surface and internal target GFP recruitment to HBpep-SA coacervates after treatment with Acetate pH 4.5 or GSH, which cause partial increases in permeability. Scale bar: 10μm. Error bars indicate standard deviation. Center of error bars indicate means. Source data are provided as a Source Data file.
